# Population genomics of the emerging forest pathogen *Neonectria neomacrospora*

**DOI:** 10.1101/2020.12.07.407155

**Authors:** Knud Nor Nielsen, Shyam Gopalakrishnan, Thorfinn Sand Korneliussen, Mikkel Skovrind, Kimmo Sirén, Bent Petersen, Thomas Sicheritz-Pontén, Iben M. Thomsen, M. Thomas P. Gilbert, Ole Kim Hansen

## Abstract

The fungal pathogen *Neonectria neomacrospora* is of increasing concern in Europe where, within the last decade, it has caused substantial damage to forest stands and ornamental trees of the genus *Abies* (Mill.). Using whole-genome sequencing of a comprehensive collection of isolates, we show the extent of three major clades within *N. neomacrospora*, which most likely diverged around the end of the last Ice Age. We find it likely that the current European epidemic of *N. neomacrospora* was founded from a population belonging to the east North American clade. All European isolates (1957-2019) had a common evolutionary history, but substantial and asymmetrical gene flow from the larger American source population could be detected. The European population shows multiple signs of having gone through a bottleneck and subsequent population expansion.

## INTRODUCTION

The decline of keystone species through encounters with exotic pests and pathogens with which they have had no long-term coevolution, is reshaping our forests. North American forests have changed within the last century as a result of the decline of American chestnut (*Castanea dentata*), elm (*Ulmus* spp.), and American Beech (*Fagus grandifolia*). In Europe, Ash (*Fraxinus* spp.) and elm (*Ulmus* spp.) have declined (Brasier and Buck, 2001; Semizer-Cuming *et al*., 2018). During the first half of the 20^th^ century, Chestnut blight (*Cryphonectria parasitica*) devastated American chestnut forests in eastern North America killing, an estimated 3.5 billion trees after its accidental introduction from Asia (Liu and Milgroom, 2007). Two pandemics within the past century caused by Dutch elm disease (*Ophiostoma spp*.) have diminished elm forests (Brasier and Buck, 2001). The introduction of the beech scale insect *Cryptococcus fagisuga* to Halifax, Canada from England around 1890, initiated an ongoing epidemic, in which the insect primes the beech trees for the subsequent infection by the fungus *Neonectria faginata* (Cale *et al*., 2017). Both beech and chestnut were important mast species in North America, and their reduction are impacting the whole ecosystem. A recent example is the Ash dieback in Europe caused by *Hymenoscyphus fraxineus*, which can likely be traced back to the introduction of as few as two strains of the pathogen from Asia (McMullan *et al*., 2018). Climate change plays a role in the movement of plants and their pathogens (Harvell *et al*., 2002), but more acute is the human-mediated movement of natural product around the world (Desprez-Loustau *et al*., 2016), and our modification of natural environments creating new opportunities for fungal pathogens (Fisher *et al*., 2012).

Fir (*Abies spp*. Mill.) constitute key tree species in the boreal forests of the northern hemisphere (Liu, 1971). In Europe, the *Abies* species with the most northern natural distribution is the European silver fir (*A. alba*), but numerous other species of various origin are widely planted in forests and landscapes throughout Northern Europe. Natural forests have been replaced over the past few centuries by monoculture plantations of exotic tree species with traits deemed desirable for human use, such as Nordmann fir (*A. nordmanniana*) that originated from around the Black Sea. It is very likely that undesirable exotic pathogens might have followed with this translocation.

Since 2008, an increasing number of reports have been published of twig blight, cankers and dieback in *Abies* sp. in northern Europe, caused by the ascomycete *Neonectria neomacrospora* (Booth & Samuels) Mantini & Samuels (anamorph *Cylindrocarpon cylindroides* var. *cylindroides* Wollenw.). According to The European and Mediterranean Plant Protection Organization (EPPO) *N. neomacrospora* was first reported in Norway in 2008, followed by Denmark in 2011, Southern Sweden in 2015, Belgium, France and England in 2017, and Finland and Germany in 2018 (EPPO, 2019). The first report of severe damage on the stand scale in Europe was in a provenance trial of *Abies lasiocarpa*, at Silkeborg, Denmark, in 2011 (Skulason *et al*., 2017). In 2013, Danish Christmas tree growers reported in a questionnaire that 86% observed damages attributed to *N. neomacrospora* (Ventzel Hansen, 2013), and awareness of the pathogen went hand in hand with the concern among growers in northern Europe who predominantly grow *Abies* spp. The apparent spread of the pathogen and the epidemic incident levels in Denmark and Norway led the EPPO’s Panel on Quarantine Pests for Forestry to add *N. neomacrospora* to its Alert list in 2017 (EPPO, 2017).

*Neonectria neomacrospora* was first described in 1910 in northern Germany (Wollenweber, 1913) and observed in western Norway in the 1940s (Robak, 1951) as well as in France, and British Columbia in the 1950s. The only previous largescale outbreak reported was from Anticosti Island, in the Gulf of St. Lawrence in Quebec described in 1965; 15 to 75% of the 40 to 50 year-old *Abies balsamea* trees were cankered. In severely affected stands, an estimated 10% of the trees had recently died. Dissection of cankers revealed that some had originated as early as 1937 (Ouellette and Bard, 1966). A strain from the Anticosti epidemic was collected along with strains from British Columbia and Norway, and compared in virulence tests on potted trees. The test showed that the Anticosti strains were significantly more aggressive, and caused more damage, than other strains (Ouellette, 1972).

In the present study, we analyse the population structure, and demographic history of *N. neomacrospora*, using whole-genome shotgun sequencing data from 71 strains sampled across the known geographical distribution of the species, including China, Europe and North America, comprising both contemporary and historical isolates. We investigate the hypothesis that the current European epidemic of *N. neomacrospora* is caused by a recent introduction of a more virulent Quebec lineage of the fungus to Europe.

## MATERIALS AND METHODS

### Collection

Since there are no prior population genetic studies of *Neonectria neomacrospora*, we aimed for as broad spatial and temporal sampling as possible. Historical sampling locations on Anticosti Island, Canada and in Norway were revisited in the contemporary sampling efforts. Five strains, collected in Norway, the Netherlands and France between 1957 and 1961, were obtained from Westerdijk Fungal Biodiversity Institute (CBS), The Netherlands and the Norwegian Institute of Bioeconomy Research, NIBIO. Five strains collected in 1967 from the outbreak centred on Anticosti Island, Quebec was obtained from The René Pomerleau Herbarium, Laurentian Forestry Centre (CFL), Canada. Two isolates from British Columbia from 1996 and 2005 were also obtained from CBS. A single *N. neomacrospora* strain from the Hubei province in China from 2014 was provided by the Herbarium Mycologicum Academiae Sinicae (HMAS). These isolates, along with isolates from Europe and Canada collected between 2015 and 2019, is listed in Table 1. All strains were sampled from individual trees, ensuring that the same individual was not sampled twice. All contemporary samples from Europe and North America have known origin, and most were geo-referenced when collected (Table 1).

**Table 1.** *Neonectria neomacrospora* isolates used in this study. Isolates marked 1 (Blue), are clones that are removed from some analyses.

| Species | Country | Location | Lat. | Long. | Year | Host | Culture collection | ENA Accession | ID | Collected or isolated by |
| --- | --- | --- | --- | --- | --- | --- | --- | --- | --- | --- |
| *N. neomacrospora* | France | Vosges | 48.0707 | 6.9509 | 1957 | Abies alba | CBS 189.61 | ERS5389223 | 001 | W. Gerlach |
| *N. neomacrospora* | Netherlands | Zwolle | 52.5112 | 6.0940 | 1961 | Abies concolor | CBS 324.61 | ERS5389224 | 004 | L. Lombard |
| *N. neomacrospora* | Belgium | Herbeumont | 49.7689 | 5.2443 | 2017 | Abies grandis | BE5104 | ERS5389225 | 018 | S. Schmitz |
| *N. neomacrospora* | Switzerland | Arboretum | 46.5102 | 6.3684 | 2017 | Abies nordmaniana |  | ERS5389226 | 019 | K.N. Nielsen |
| *N. neomacrospora* | Norway | Os | 62.4965 | 11.2233 | 1958 | Abies alba |  | ERS5389227 | 005 | Robak |
| *N. neomacrospora* | Norway | Fana | 60.2741 | 5.3954 | 1961 | Abies alba | CBS 503.67 | ERS5389228 | 049 | R. Roll-Hansen |
| *N. neomacrospora* | Norway | Fana | 60.2716 | 5.3866 | 1961 | Abies alba | NO 61-62/1 | ERS5389229 | 051 | R. Roll-Hansen |
| *N. neomacrospora* | Norway | Fana | 60.2600 | 5.3400 | 2019 | Abies lasiocarpa | NO 252125 | ERS5394065 | 093 | J.-O. Skage |
| *N. neomacrospora* | Norway | Fana | 60.2600 | 5.3400 | 2019 | Abies lasiocarpa | NO 252130 | ERS5389230 | 095 | J.-O. Skage |
| *N. neomacrospora* | Norway | Fana | 60.2600 | 5.3400 | 2019 | Abies lasiocarpa | NO 252140 | ERS5389231 | 097 | J.-O. Skage |
| *N. neomacrospora* | Denmark | Arboretum <sup>1</sup> | 55.8691 | 12.5033 | 2015 | Abies fargesii |  | ERS5389232 | 020 | K.N. Nielsen |
| *N. neomacrospora* | Denmark | Arboretum <sup>1</sup> | 55,8667 | 12,5097 | 2015 | Abies lasiocarpa |  | ERS5389233 | 021 | K.N. Nielsen |
| *N. neomacrospora* | Denmark | Arboretum | 55.8642 | 12.5119 | 2016 | Abies lasiocarpa |  | ERS5389234 | 022 | K.N. Nielsen |
| *N. neomacrospora* | Denmark | Arboretum <sup>1</sup> | 55,8648 | 12,5117 | 2016 | Abies lasiocarpa |  | ERS5389235 | 023 | K.N. Nielsen |
| *N. neomacrospora* | Denmark | Arboretum | 55.8642 | 12.5093 | 2015 | Abies pinsapo |  | ERS5389236 | 025 | K.N. Nielsen |
| *N. neomacrospora* | Denmark | Arboretum <sup>1</sup> | 55,8649 | 12,5107 | 2016 | Abies chensiensis |  | ERS5389237 | 026 | K.N. Nielsen |
| *N. neomacrospora* | Denmark | Arboretum <sup>1</sup> | 55,8673 | 12,5096 | 2016 | Abies procera |  | ERS5389238 | 027 | K.N. Nielsen |
| *N. neomacrospora* | Denmark | Silkeborg | 56.1634 | 9.5745 | 2015 | Abies nordmaniana |  |  | ref | K.N. Nielsen |
| *N. neomacrospora* | Denmark | Silkeborg | 56.1627 | 9.5750 | 2016 | Abies nordmaniana |  | ERS5389239 | 029 | K.N. Nielsen |
| *N. neomacrospora* | Denmark | Silkeborg | 56.1632 | 9.5757 | 2016 | Abies nordmaniana |  | ERS5389240 | 031 | K.N. Nielsen |
| *N. neomacrospora* | Denmark | Silkeborg | 56.1625 | 9.5745 | 2016 | Abies nordmaniana |  | ERS5389241 | 032 | K.N. Nielsen |
| *N. neomacrospora* | Denmark | Silkeborg <sup>2</sup> | 56.1626 | 9.5741 | 2016 | Abies nordmaniana |  | ERS5389242 | 033 | K.N. Nielsen |
| *N. neomacrospora* | Denmark | Silkeborg | 56.1626 | 9.5718 | 2016 | Abies nordmaniana |  | ERS5389243 | 035 | K.N. Nielsen |
| *N. neomacrospora* | Denmark | Silkeborg <sup>2</sup> | 56,1632 | 9,5739 | 2016 | Abies nordmaniana |  | ERS5389244 | 036 | K.N. Nielsen |
| *N. neomacrospora* | Denmark | Thy | 57.0242 | 8.5987 | 2015 | Abies nordmaniana |  | ERS5389245 | 037 | K.N. Nielsen |
| *N. neomacrospora* | Denmark | Thy | 57.0241 | 8.5989 | 2015 | Abies nordmaniana |  | ERS5389246 | 038 | K.N. Nielsen |
| *N. neomacrospora* | Denmark | Christiansfeld <sup>3</sup> | 55.3643 | 9.4378 | 2018 | Abies procera |  | ERS5389247 | 039 | K.N. Nielsen |
| *N. neomacrospora* | Denmark | Christiansfeld | 55.3639 | 9.4378 | 2018 | Abies procera |  | ERS5389248 | 040 | K.N. Nielsen |
| *N. neomacrospora* | Denmark | Christiansfeld | 55.3637 | 9.4379 | 2018 | Abies procera |  | ERS5389249 | 041 | K.N. Nielsen |
| *N. neomacrospora* | Denmark | Christiansfeld <sup>3</sup> | 55,3632 | 9,4378 | 2018 | Abies procera |  | ERS5389250 | 042 | K.N. Nielsen |
| *N. neomacrospora* | Denmark | Christiansfeld | 55.3630 | 9.4378 | 2018 | Abies procera |  | ERS5389251 | 043 | K.N. Nielsen |
| *N. neomacrospora* | Denmark | Christiansfeld | 55.3626 | 9.4377 | 2018 | Abies procera |  | ERS5389252 | 044 | K.N. Nielsen |
| *N. neomacrospora* | Denmark | Christiansfeld | 55.3625 | 9.4378 | 2018 | Abies procera |  | ERS5389253 | 045 | K.N. Nielsen |
| *N. neomacrospora* | Denmark | Christiansfeld | 55.3621 | 9.4376 | 2018 | Abies procera |  | ERS5389254 | 046 | K.N. Nielsen |
| *N. neomacrospora* | Denmark | Bommerlund | 54.8790 | 9.3447 | 2018 | Abies nordmaniana |  | ERS5389255 | 081 | K.N. Nielsen |
| *N. neomacrospora* | Denmark | Bommerlund | 54.8782 | 9.3442 | 2018 | Abies nordmaniana |  | ERS5389256 | 082 | K.N. Nielsen |
| *N. neomacrospora* | Denmark | Skelhusmarken | 56.7781 | 9.8417 | 2015 | Abies nordmaniana |  | ERS5389257 | 103 | K.N. Nielsen |
| *N. neomacrospora* | Denmark | Skelhusmarken | 56.7789 | 9.8423 | 2015 | Abies nordmaniana |  | ERS5389258 | 104 | K.N. Nielsen |
| *N. neomacrospora* | Denmark | Skelhusmarken | 56.7790 | 9.8420 | 2015 | Abies nordmaniana |  | ERS5389259 | 105 | K.N. Nielsen |
| *N. neomacrospora* | Denmark | Varde | 55.5957 | 8.5284 | 2016 | Abies grandis |  | ERS5389260 | 107 | K.N. Nielsen |
| *N. neomacrospora* | Denmark | Varde | 55.5880 | 8.5235 | 2016 | Abies grandis |  | ERS5389261 | 108 | K.N. Nielsen |
| *N. neomacrospora* | Finland | Mustila | 60.7315 | 26.4214 | 2018 | Abies sp. |  | ERS5389262 | 048 | A. Uimari |
| *N. neomacrospora* | Finland | Jarvenpaa | 60.4664 | 25.0896 | 2019 | Abies sp. |  | ERS5389263 | 084 | A. Uimari |
| *N. neomacrospora* | Finland |  | 60.1919 | 24.9368 | 2019 | Abies sp. |  | ERS5389264 | 085 | A. Uimari |
| *N. neomacrospora* | Finland | Espoo L 2 | 60.2014 | 24.8041 | 2019 | Abies sp. |  | ERS5389265 | 086 | A. Uimari |
| *N. neomacrospora* | Finland | Salo 15 | 60.3841 | 23.0868 | 2019 | Abies sp. |  | ERS5389266 | 088 | A. Uimari |
| *N. neomacrospora* | Wales | Llanbrynmair | 52.5143 | -3.6887 | 2016 | Abies procera |  | ERS5389267 | 070 | J. McMinn |
| *N. neomacrospora* | Wales | Llanbrynmair | 52.4674 | -3.6808 | 2016 | Abies sp. |  | ERS5389268 | 072 | J. McMinn |
| *N. neomacrospora* | England | Kent | 51.0762 | 0.4635 | 2017 | Abies nordmaniana |  | ERS5389269 | 078 | J. McMinn |
| *N. neomacrospora* | England | Kent | 51.0761 | 0.4535 | 2017 | Abies sp. |  | ERS5389270 | 080 | J. McMinn |
| *N. neomacrospora* | Canada | Anti-costi | 49.8200 | -64.3500 | 1967 | Abies balsamea | CFL 961 | ERS5389271 | 010 | P.O. Duguay |
| *N. neomacrospora* | Canada | Anti-costi | 49.8551 | -64.1037 | 1967 | Abies balsamea | CFL 962 | ERS5389272 | 011 | P.O. Duguay |
| *N. neomacrospora* | Canada | Anti-costi | 49.8551 | -64.1037 | 1967 | Abies balsamea | CFL 965 | ERS5389273 | 012 | P.O. Duguay |
| *N. neomacrospora* | Canada | Anti-costi | 49.7493 | -63.1044 | 2018 | Abies balsamea |  | ERS5389274 | 052 | K.N. Nielsen |
| *N. neomacrospora* | Canada | Anti-costi | 49.8225 | -63.8326 | 2018 | Abies balsamea |  | ERS5389275 | 053 | K.N. Nielsen |
| *N. neomacrospora* | Canada | Anti-costi | 49.7564 | -63.0759 | 2018 | Abies balsamea |  | ERS5389276 | 054 | K.N. Nielsen |
| *N. neomacrospora* | Canada | Anti-costi | 49.8266 | -63.8492 | 2018 | Abies balsamea |  | ERS5389277 | 055 | K.N. Nielsen |
| *N. neomacrospora* | Canada | Anti-costi | 49.5589 | -62.6887 | 2018 | Abies balsamea |  | ERS5389278 | 056 | K.N. Nielsen |
| *N. neomacrospora* | Canada | Anti-costi | 49.7563 | -63.0761 | 2018 | Abies balsamea |  | ERS5389279 | 058 | K.N. Nielsen |
| *N. neomacrospora* | Canada | Anti-costi | 49.8224 | -63.8323 | 2018 | Abies balsamea |  | ERS5389280 | 059 | K.N. Nielsen |
| *N. neomacrospora* | Canada | Anti-costi | 49.9146 | -64.3410 | 2018 | Abies balsamea |  | ERS5389281 | 060 | K.N. Nielsen |
| *N. neomacrospora* | Canada | Anti-costi | 49.9150 | -64.3394 | 2018 | Abies balsamea |  | ERS5389282 | 061 | K.N. Nielsen |
| *N. neomacrospora* | Canada | Port Cartier | 50.0300 | -66.8700 | 1967 | Abies balsamea | CFL 963 | ERS5389283 | 015 |  |
| *N. neomacrospora* | Canada | Baie-Comeau | 49.2300 | -68.1600 | 1967 | Abies balsamea | CFL 964 | ERS5389284 | 016 |  |
| *N. neomacrospora* | Canada | Cote Nord | 50.2998 | -66.4364 | 2018 | Abies balsamea |  | ERS5389285 | 062 | K.N. Nielsen |
| *N. neomacrospora* | Canada | British | 52.0000 | -126.0000 | 1996 | Tsuga heterophylla | CBS 118985 | ERS5389286 | 007 | S. Shamoun |
| *N. neomacrospora* | Canada | Cobble Hill | 48.7002 | -123.5912 | 2018 | Abies lasiocarpa |  | ERS5389287 | 063 | K.N. Nielsen |
| *N. neomacrospora* | Canada | Cobble Hill | 48.7005 | -123.5911 | 2018 | Abies lasiocarpa |  | ERS5389288 | 064 | K.N. Nielsen |
| *N. neomacrospora* | Canada | Cobble Hill | 48.7004 | -122.5910 | 2018 | Abies lasiocarpa |  | ERS5389289 | 065 | K.N. Nielsen |
| *N. neomacrospora* | Canada | Cobble Hill | 48.7002 | -123.5910 | 2018 | Abies lasiocarpa |  | ERS5389290 | 066 | K.N. Nielsen |
| *N. neomacrospora* | Canada | Vancouver Island | 49.3471 | -124.6259 | 2005 | Arceuthobium tsugensw. on Abies balsamea | CBS 118984 | ERS5389291 | 106 | L. Reitman |
| *N. neomacrospora* | China | Shennongjia | 31.7300 | 110.6800 | 2014 | Pinus sp. | HMAS 252906 | ERS5389292 | 017 | Zhao-Qing Zeng |

### Isolating pure cultures

Macroconidia were collected from sporodochia on the bark of infected *Abies* sp., using the tip of a needle. When sporodochia were not available, the fungus was isolated from the wood and microconidia were collected from these cultures. Axenic single-spore cultures were derived by plating a small number of conidia diluted in water on potato dextrose agar (PDA) plates, which allowed conidia to separate. After 24 h of incubation, plates were observed under a dissection microscope at 50× magnification and single germinating conidia were collected and transferred to new PDA plates. Single-spore cultures were maintained in 20 % (v/v) glycerol at −80 °C.

### DNA extraction and sequencing

Isolates were transferred to potato dextrose broth (PDB) for 4-5 days at room temperature, and the mycelium was collected on Whatman filter paper (grade 1), rinsed with water and lyophilised. 20-40 mg dried mycelium was homogenised with 200 mg 1 mm zirconia beads in a bead mill (Retsh Mixer Mill MM301) prior to DNA extraction. DNA was extracted with the DNeasy UltraClean® Microbial DNA Isolation Kit (Qiagen) with the addition of Proteinase K 1% to the lysis mix, and a prolonged lysis incubation of 2 hours at 62 °C. DNA was purified with the DNeasy PowerClean Pro Cleanup Kit, and concentrations were determined using a Qubit 3 Fluorometer with the Qubit™ dsDNA BR Assay Kit.

DNA extracts were fragmented by sonication to 200-800 bp using the Covaris M220. Illumina compatible sequencing libraries were constructed following the BEST protocol described in Carøe *et al*. (2018), using 100-300 ng dsDNA, and dual-indexed with seven bp indexes. Extraction, library and index PCR blanks were included to evaluate for potential contamination during the library building process. No blanks amplified in the qPCR quantification step, and thus the blanks were therefore not sequenced. To ensure library complexity, amplification was done in duplicates and subsequently pooled prior to purification with SPRI-beads. Indexed libraries were quantified on a 5200 Fragment Analyzer System (Agilent), and an equimolar pool of all libraries was produced. The pooled library was purification using a BluePippin (Sage Science, Beverly, MA, USA), selecting fragments between 200 bp and 1000 bp. Libraries were sequenced on one lane of an Illumina NovaSeq 6000 SP 150 PE sequencing, at the Danish National High-Throughput DNA Sequencing Centre.

### Trimming and adapter removal

Reads were trimmed, removing Illumina adapter and primer sequences and bases at read ends with Phred quality below 20 (-q20), while only keeping reads longer than 80 bp. This was performed using AdapterRemoval (v.2.2.4)(Schubert, Lindgreen and Orlando, 2016), options: [--trimns --trimqualities --minquality 20 --minlength 80].

### Genome assembly and gene prediction

Trimmed reads were de-novo assembled using SPAdes v.3.13.1 (Bankevich *et al*., 2012) (kmers 21, 33, 55, 77, 99, 127) using mismatch and short indel correction with the Burrow-Wheeler Aligner, BWA-MEM v.0.7.16a (Li, 2013). Assemblies were improved using Pilon v.1.22 (Walker *et al*., 2014). The assembly summary statistics were calculated using Quast v5.0.2 (Mikheenko *et al*., 2018) (Table S1).

Gene prediction on the polished assemblies was performed using the Funannotate pipeline v. 1.6.0, (see URLs), utilising two gene prediction tools: AUGUSTUS (Stanke and Morgenstern, 2005) and GeneMark-ES (Besemer and Borodovsky, 2005), with *Fusarium graminearum* as a model for the AUGUSTUS gene predicter and BAKER1 (Hoff *et al*., 2016) for the training of GeneMark-ES. Consensus gene models were found with EvidenceModeler (Haas *et al*., 2008).

### Mating types

The mating type of each isolate was identified in the genome assemblies using the NCBI BLAST+ v2.10.0, with a blast database build on the nucleotide sequences of the two *N. neomacrospora* mating type genes MAT1.1.1 and MAT1.2.1, with the GeneBank assessions: MT457585.1 and MT457570.1 (Stauder *et al*., 2020).

### Reads mapping, variant calling and filtering

Three variant dataset were generated: 1. A set of 28 thousand bi-allelic, single nucleotide polymorphisms (SNPs) with a minimum sequencing depth of 5 in 80 % of the samples, used for linkage disequilibrium (LD) analysis; 2. A subset of 8905 SNPs with a minimum distance of two kb, used for PCA and Admixture analysis. These two sets were generated as follows: For each isolate, the reads were mapped to the *N. neomacrospora* strain KNNDK1 reference genome (unpublished), with BWA-MEM v.0.7.16a, using default parameters. Duplicate reads were marked, reads were realigned for short indels and variants were called with GATK v.4.1.2.0, with ‘-ERC GVCF’ cohort analysis workflow mode and ploidy set to 1. The GATK module ‘VariantFiltration’ was used to quality filter SNVs based on the values ‘QUAL < 30.0’, ‘QD < 25.0’, ‘SOR > 3.0’, ‘FS > 10.0’, ‘MQ < 55.0’, ‘MQRankSum < −0.4’ and ‘ReadPosRankSum < −2.0’. SNPs were hard-filtered using VCFtools v.0.1.16 (Danecek *et al*., 2011) to only include bi-allelic SNPs that had a minimum per sample sequencing depth of five (disregarding duplicates) and was sequenced in a minimum 80 % of the strains. No evidence of chromosomal aneuploidy has been found (Figure S1), and ploidy was therefore set to 1.

The third dataset used for estimating the population scaled mutation rate (θ), were called using BCFtools (1.9-94-g9589876) (Li *et al*., 2009). This was done by using a combination of BCFtools mpileup and call (--ploidy 1) using a mapping quality filter of 30 and a basequality filter of 20 together with default parameters including BAQ (Li, 2011).

### Population structure from PCA and Admixture

We analysed the population structure of 71 *N. neomacrospora* isolates using two methods: Principal-component analysis (PCA) using SNPRelate v1.18.1 (Zheng *et al*., 2012), and Admixture v1.3.0 (Alexander, Novembre and Lange, 2009). For the admixture analysis data was clone-censured using the R package poppr (Kamvar, Tabima and Grünwald, 2014), removing six isolates from three clones for in Denmark. The admixture analysis was run 200 times of each K, and the clustering with the lowest cross-validation error for each K was visualized. Both analyses were visualised with ggplot2 v3.2.1 (Wickhm, 2019) in R v3.6.1 (R Core Team, 2019).

### Linkage disequilibrium

The level of linkage disequilibrium (LD) in the European and Quebec populations was calculated as pairwise r^2^ within 50 kb windows between all SNPs using PLINK v.1.90b3o (see URLs). Distances between SNPs were calculated and SNPs aggregated in distance bins of 100 bp for subsequent calculation of mean and sd for the calculated r^2^ values. The LD decay plot was made with the R package ggplot2.

### Estimates of the population scaled mutation rate (θ), neutrality test statistics and population differentiation

Different estimators of the population scaled mutation rate (θ) has been proposed and take the general form for a locus with S sites and n chromosomes: 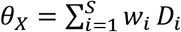, here Di denotes the number of derived alleles for site *i* with *w*_*i*_ being different ‘weights’ given by the number of derived alleles. The classic Watterson estimator is then written as 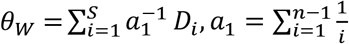. In this case all weights are the same across all categories of derived alleles, this is different from the pairwise estimator of theta which has the highest weights on the intermediate categories 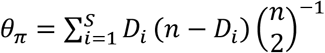. These two estimators do not use information about the polarisation of the outgroup in contrast to the Fay & Wu estimator: 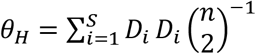 (Fay and Wu, 2000). For the sake of completeness, we have also included Fu and Li’s L theta estimate which is simply given by the singleton category 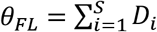, *f or D*_*i*_ = 1 (Fu and Li, 1993). These are all unbiased estimators of the same quantity and any difference between these estimators can be used as a test statistic for finding deviations from neutrality (Durrett, 2008; Achaz, 2009), the most widely used being Tajima’s D (θπ -θW) (Tajima, 1989). We used BCFtools (1.9-94-g9589876) to call (haploid) genotypes and used custom R scripts (see github repository), for estimating per-site thetas and performing the window based neutrality test using 5kb windows, due to the difference in effective number of sites between windows we discarded those windows that had less than half of the average number of sites for each chromosome.

Sample size bias in Tajima’s D was investigated by rarefaction of the European population to n = 15, the same size as the Quebec sample. Based on the variant dataset 2, Tajima’s D was calculated on 100 random subsamples using the [--max-indv] option in VCFtools. Mean values for the 100 iterations of each 10 kb window across the genome was calculated. Population differentiation was estimated by calculating pairwise F_ST_ fixation indices among populations (Wright, 1951), this was done by using the moment estimator (Weir and Cockerham, 1984).

Data used for these analyses was clone-censured, excluding all but one isolate of each of the three clones sampled in Denmark, reducing the total sample size from 71 to 65 isolates.

### Phylogeny

Predicted protein data was used for the identification of orthologous gene families. The protein transcripts of 1418 single-copy orthologous gene clusters were aligned using MAFFT v. 7.402 [option: linsi] (Katoh and Standley, 2013). Aligned genes were filtered on amount of gaps and inter-gene distance, leaving 51 genes with less than two per cent gaps and a minimum inter-gene distance of 10kb (on the reference genome KNNDK1). Substitution models for each codon position in each gene were predicted using ModelFinder (Kalyaanamoorthy *et al*., 2017) as implemented in IQtree v.1.6.12, and used with the concatenated protein alignment to generate a consensus maximum likelihood phylogeny based on 100 trees. The consensus tree was subsequently validated with 100 bootstrap replicates using IQtree v.1.6.12 (Nguyen *et al*., 2015; Chernomor, von Haeseler and Minh, 2016). The outgroup *N. major*, is not shown in the phylogeny (Figure 2, S3-S5).

Divergence time analysis was performed applying a Coalescent Constant Population model in BEAST v2.6.1 (Bouckaert *et al*., 2014) suited for single-species studies, and a strict clock rate under the assumption that there is very little rate heterogeneity within *N. neomacrospora*. Only the third codon position was used for calculating the time to the most recent common ancestor (TMRCA), to reduce the effect of purifying selection on time estimates. The third codon positions of the 51 genes were run as six partitions, based on the merger by ModelFinder. All partitions were run with the HKY substitution model. We used linked trees, linked clocks and unlinked site models with estimated substitution rates. The Markov chain Monte Carlo (MCMC) was run with 100 million steps storing every 5000 steps. Effective sample size (ESS) were inspected using Tracer 1.7 (Rambaut *et al*., 2018); all ESS values were above 950 and considered converged. Posterior probabilities of these trees were summarized using the maximum clade credibility method implemented in TreeAnnnotator v2.6.0 from the Beast2 package (Bouckaert *et al*., 2019) and [option: 10% burnin; median heights], and plotted using FigTree v1.4.4 (see URLs).

Mitochondrial genomes were assembled by read-mapping to the mitochondrial reference genome KNNDK1. Reads were aligned to reference with BWA-MEM v.0.7.16a, the Samtools v. 1.9 (Li *et al*., 2009) [--dedup] option was used to remove duplicated reads, and angsd v.0.929 (Korneliussen, Albrechtsen and Nielsen, 2014) [--doFasta2 -setMinDepth 20] called the most common base for generating fasta assemblies where bam coverage was >20x. Mitochondrial genomes were aligned with MAFFT v. 7.402 with the local alignment option [-linsi] for high accuracy.

The substitution models best fitting the mitochondrial data were selected using ModelFinder (Kalyaanamoorthy *et al*., 2017) as implemented in IQtree (Nguyen *et al*., 2015; Chernomor, von Haeseler and Minh, 2016). A maximum likelihood consensus tree was made with IQtree using 100 bootstrap replicates.

To identify unique haplotypes and visualize the number of substitutions separating them, we constructed a median spanning network using POPART v1.7 (Leigh and Bryant, 2015). The analysis was based on the full mitochondrial genome alignment described above.

### Demographic reconstruction

The Extended Bayesian skyline plot (EBSP) implemented in BEAST v2.6.1 (Bouckaert *et al*., 2014) was used to infer demographic history. The analysis was conducted with the 51 single-copy genes selected for the nuclear phylogeny. Only the third codon positions were used to minimize the effects of selection on time estimates of recent evolutionary events. All partitions were run with a HKY substitution model, with gamma site heterogeneity and six categories, under the assumption of a strict clock rate. The inference was calibrated using tip-dates for all strains. The Markov Chain Monte Carlo (MCMC) analyses were first performed with short runs with a chain length of 10^6^ to optimize the scale factors of the priors. The analysis was then run for 10^8^ generations, sampling every 1000th iteration after an initial burn-in of 10%. The performance of the MCMC process was checked for stationarity and large effective sample sizes in Tracer. The skyline was calculated and plotted using the plotEBSP R script available at the BEAST2 web site (see URLs).

Current and ancestral population sizes were estimated for the European and the Quebec populations, as were migration rates between the two populations determined using the python package moments (Jouganous *et al*., 2017), that uses a diffusion approximation for identifying the demographic parameters from the estimated site frequency spectrum (SFS). The 2-d SFS (two population SFS) between the Quebec and European populations was estimated using angsd v.0.931. Using the estimated SFS, we fitted four demographic models: following the split of the two population we model an asymmetric migration between Europe and Quebec and either: 1. Population growth in both populations, 2. growth only in QC, 3. growth only in EU, or 4. a constant population size in both populations (i.e. no growth). The different models were compared using the log likelihood of the estimated parameters under the model.

## RESULTS

We sequenced the whole genomes of 71 *N. neomacrospora* strains collected from Europe (n=49), North America (n=21) and China (n=1), spanning from 1957 to 2019. All samples were collected from *Abies* spp., except the Chinese strain, which is reported to originate from a *Pinus* sp. Strains were sequenced to a mean 30 fold coverage across the nuclear genome (Table S1).

### Population structure by PCA

We observed no replacement of the old European population of *N. neomacrospora* by a different lineage. All samples from China, British Columbia, Quebec and Europe clustered into lineages that reflect the geographical origin of sampling. Historic samples clustered with the contemporary samples of their respective geographic sampling areas (Figure 1ab), and this temporal stratification did not reveal a translocation of strains within the last 50 years. No isolates show inter-population placements within the PCA, reflecting intermediates genotypes. If hybridization and introgression are present, they could not be detected despite the 28 thousand SNPs analysed.

**Figure 1.**
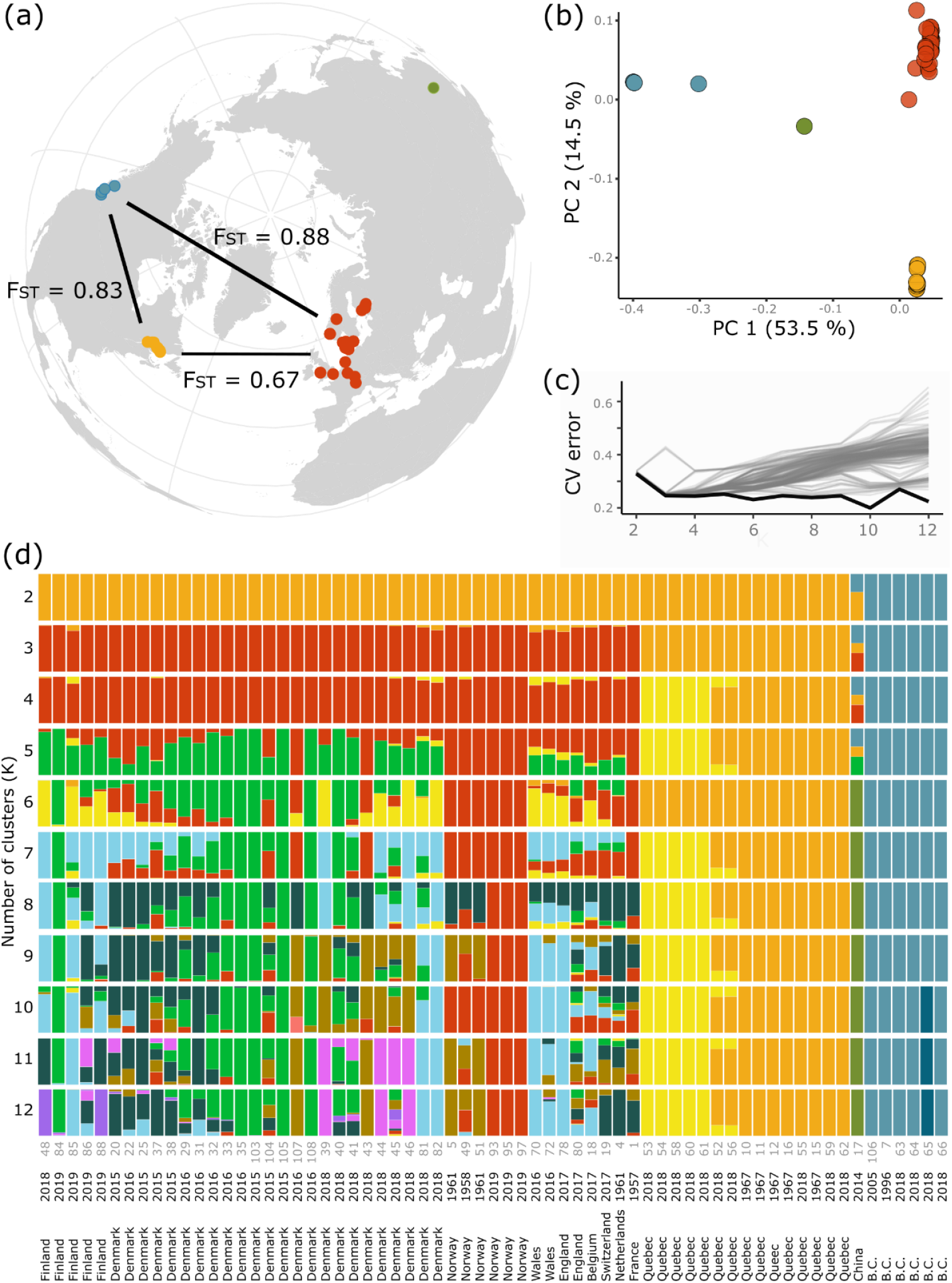
F_ST_, Principal component analysis and Admixture analysis of the sequenced strains of *Neonectria neomacrospora* based on 8905 bi-allelic SNPs. a) F_ST_ values between the three populations. b) PC 1 of the PCA describe 53.5% of the variation in the data, separates the British Columbia and China from Europe and Quebec. EU and QC are separated by PC 2. Historical samples within EU and QC are within the dashed circles. Admixture was run 200 of K 1-12, c) Shows the cross-validation error associated with each value of K, the optimal clustering of each K (bottom line) is shown in d). d) gives the estimated likely ancestral clusters given a clustering into K groups.

### Admixture

The ADMIXTURE analysis found that the K-value (number of inferred ancestral populations) with the lowest cross-validation error was ten (Figure 1c). This clustering corresponded with clustering to the geographical origin of the strains, i.e. British Columbia, Quebec, Europe and China, as well as substantial subclustering of within Europe (Figure 1d). The cross-validation error is similar comparing K values between three and nine. At K=4, the Quebec samples were split into two clusters, the minor group all originated of the Anticosti Island and were all collected in 2018. Twelve European samples show a partly shared ancestry with this minor Quebec group at K=4. This could be a signal of introgression from Quebec into the European population, and is seen for a variable number of isolates for all K between four and 11.

### Mating type

Disregarding the one sample Chinese admixture cluster; both mating-type MAT1.1.1 and MAT1.2.1 were found in all ancestral groups identified in the admixture analysis where K equals eight or less. This means that both mating types were present in all sampled regions, which is in line with expectations based on frequent observations of the sexually produced perithecia. The mating-type MAT1.1.1 were the most frequently sampled of the two with, n_MAT1.1.1_=39 compared to n_MAT1.2.1_=37. Clone-correction removes both mating types bringing the counts down to: n_MAT1.1.1_ =32 and n_MAT1.2.1_=28, respectively.

### Nuclear phylogeny

For the nuclear phylogeny, a genome-wide selection of 51 single copy ortholog genes was used, partitioned into the three different codon positions per gene. ModelFinder merged the 153 subsets into 16 and assigned the best fitting substitutions models. While the maximum likelihood phylogeny was made from this dataset, the MCMC phylogeny was only based on the third codon position, corresponding to six partitions.

The bootstrap analyses on the maximum-likelihood consensus phylogeny (Figure S3) and the Bayesian MCMC phylogeny (Figure S4) were concurrently giving 100% support for a split into four monophyletic clades matching the sampling regions, Europe, Quebec, China and British Colombia (Figure 2). Where the PCA and admixture analyses had the Chinese lineage as an intermediate between British Columbian and European genotypes, it is clear from the phylogeny that the *N. neomacrospora* consist of at least three major clades represented by the British Columbian, the Chinese and the combined Europe-Quebec lineages.

**Figure 2.**
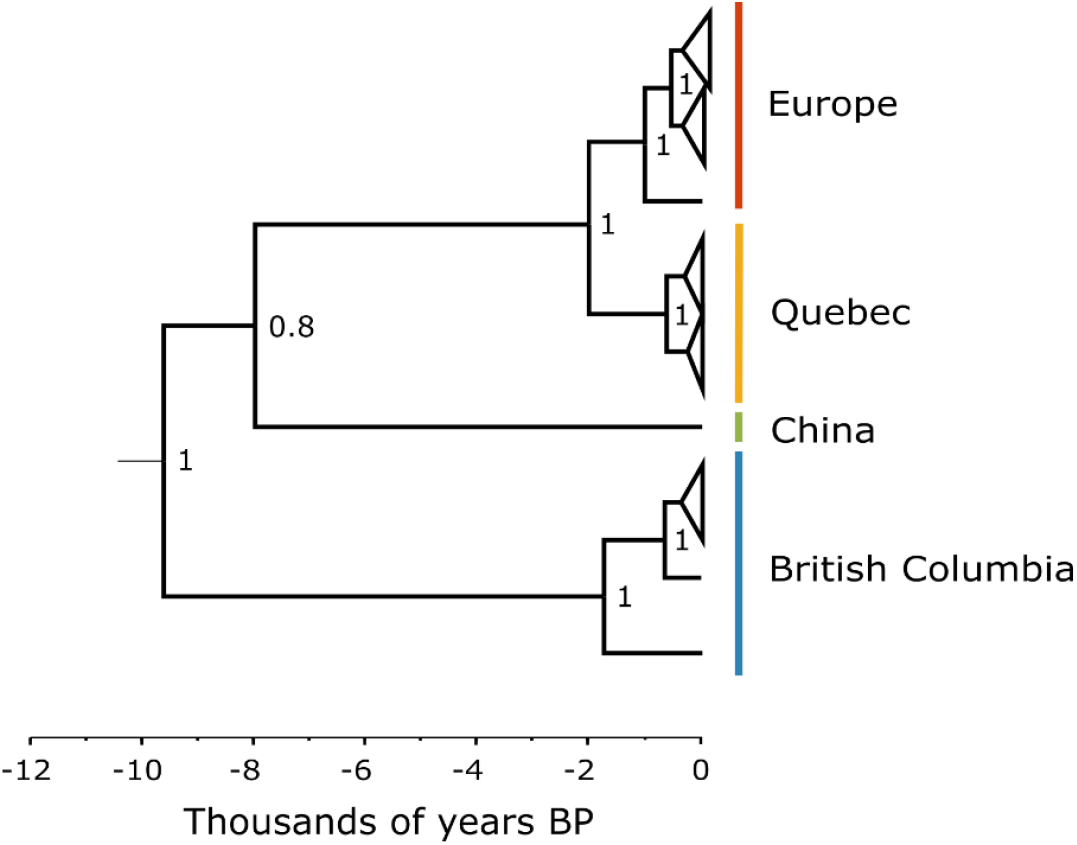
Evolutionary history of *Neonectria neomacrospora*. The tree topology is supported by both Bayesian and frequency-based phylogenies. Bayesian inference: Node labels show the posterior probability of splits (range: 0-1). Splits are set at median tree height, given by the posterior density of the split age. The corresponding maximum likelihood consensus tree gives bootstrap values of 100 to the four monophyletic clades corresponding to the four regions: Europe, Quebec, China and British Columbia.

### Mitochondrial haplotype network

The haplotype network included 218 informative sites forming 23 haplotypes, representing between one and twenty-four isolates (Figure 3). The largest intra-population haplotype divergence is found within the Quebec population with a distance of 15 nucleotide differences, more than three times the maximum distance found within the European population (four nucleotide difference). The two major European haplotypes, including 24 and 8 isolates respectively, do not correspond to the large groups identified in the phylogeny from 51 nuclear genes. The two groups are not geographically structured either. The two Quebec groups identified in the admixture analyses (K=4-5,7-12) on nuclear genome SNPs correspond to splitting the Quebec haplotypes into two groups: one with the four haplotypes closest to the BC haplotypes and a minor group containing the remaining three haplotypes (five isolates) (Figure 3).

**Figure 3.**
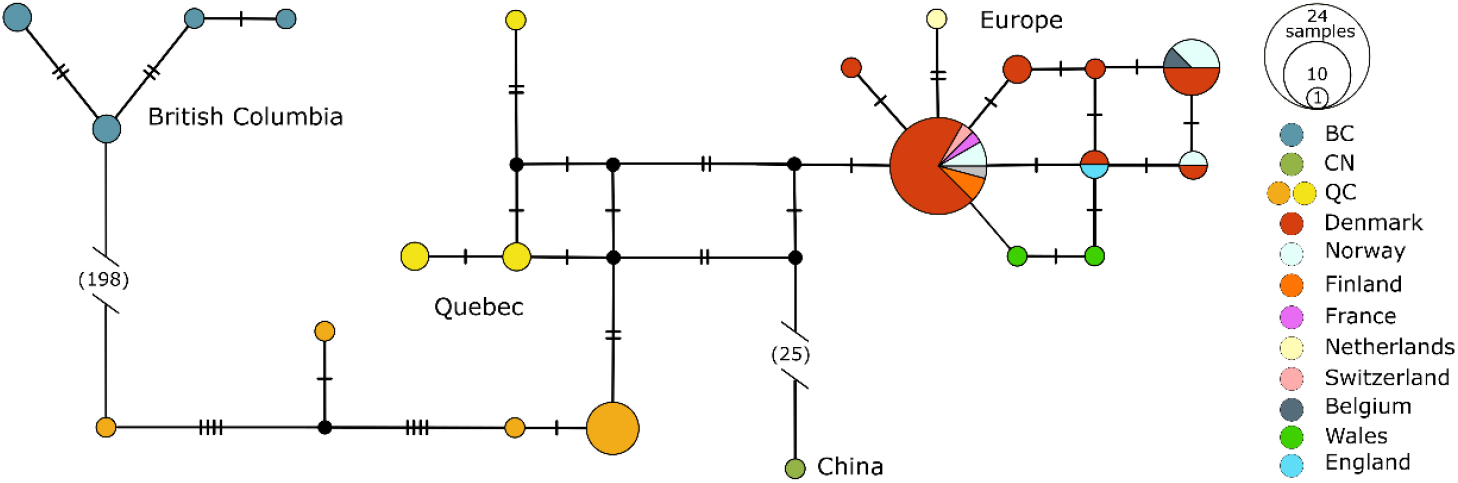
Haplotype network of *Neonectria neomacrospora* mitogenomes. Median-spanning network of 23 mitogenome haplotypes found across 65 clone-censured isolates. Each haplotype is represented by a circle, and the circle size indicates the relative frequency of haplotype. Circles are coloured according to their sample sites. QC is given two colours corresponding to the two cluster identified in the admixture analysis on nuclear SNPs. Black dots indicate haplotypes not present in the data. Hatches and numbers in brackets indicate the number of nucleotide differences between haplotypes.

### Theta estimates and neutrality test statistics

The overall estimates of theta are θ_pi_=32109.87 and θ_W=_36575.04 for the entire genome. See Figure 4 for the local Watterson and pairwise theta estimates for all three populations estimated for 5 kb regions across the genome. The diversity found in the European population is higher than the observed diversity in the two North American populations sampled, both in the number of variable sites θ_W_ and in the pairwise diversity measure θ_T_ (Table 2a). This is in contrast to the pattern observed in the mitochondrial haplotype network (Figure 3).

**Table 2(a).** Summary statistic incl estimators of theta.

| Origin | n | Sites | Variable sites | $\theta_w$ | $\theta_\pi$ | $\theta_H$ | $\theta_L$ |
| --- | --- | --- | --- | --- | --- | --- | --- |
| Europe | 43 | 23,912,088 | 127,590 | 0.0012 | 0.0011 | 0.0030 | 0.0012 |
| Quebec | 15 | 23,837,969 | 70,091 | 0.0009 | 0.0008 | 0.0010 | 0.0010 |
| BC | 6 | 23,554,712 | 83,554 | 0.0016 | 0.0014 | 0.0020 | 0.0019 |

For the three populations we found the number of sites with data (Sites) along with the number of variable sites (Variable sites) defined as having atleast two different alleles. We also applied four different estimators of theta (Watterson, Pairwise, Fay and Wu's H and Fu's L). The theta estimates were estimates across the entire locus (all nuclear chromosomes) and rescaled to a per site estimate. We remark that Fay and Wu's H and Fu's L requires proper polarization to the ancestral states whereas the Watterson and the pairwise is invariant with regards to the choice of polarization.

**Figure 4.**
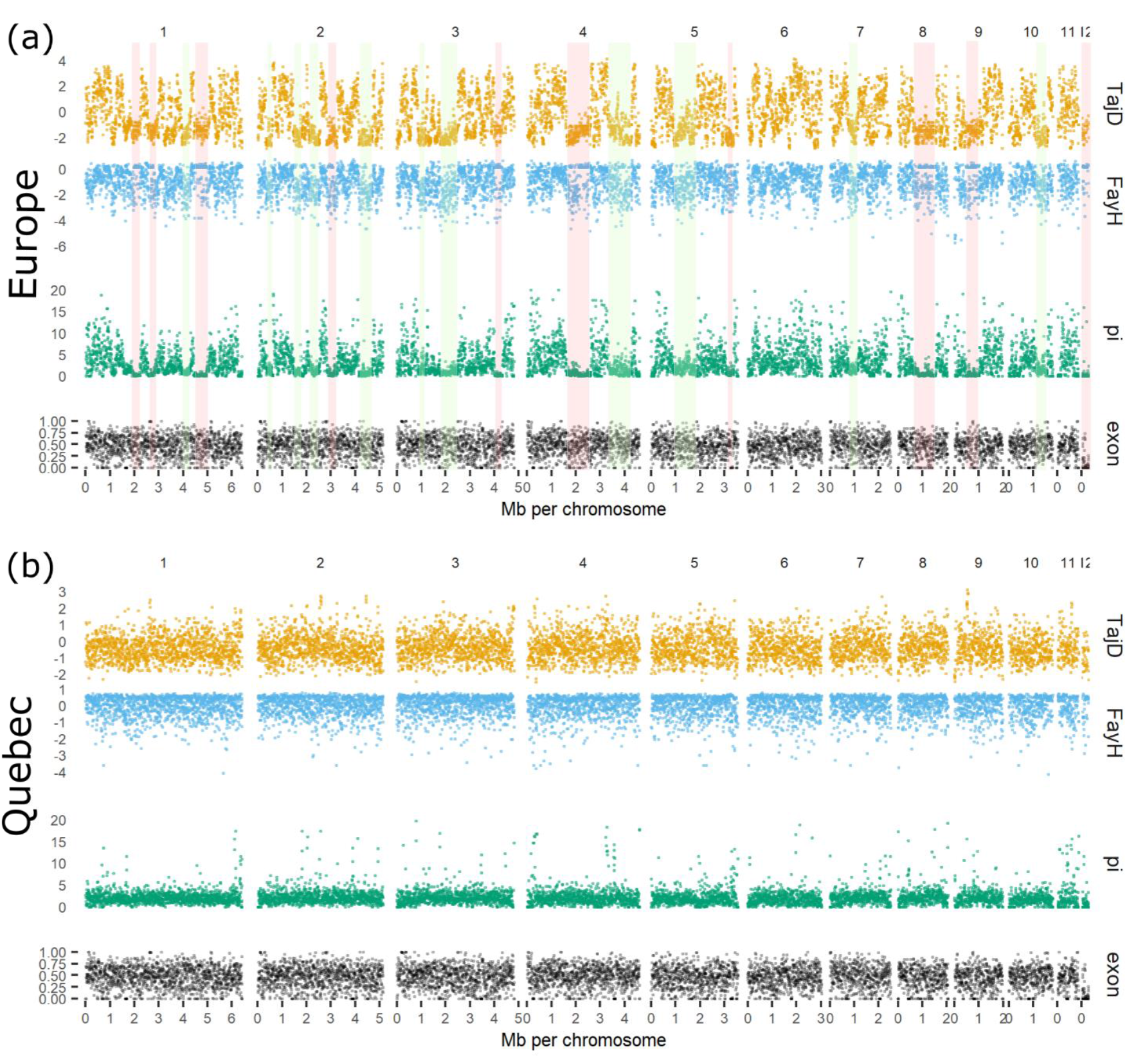
Window based statistic with 5 kb windows across the genome of *Neonectria neomacrospora*. The two panel each gives local estimates of theta pi and neutrality test statistic for Tajima’s D and Fay’s H, as well as the exon coverage in fractions of the windows. Subpanel A) summarises values for the 49 European isolate, where subpanel B) summarises the 15 isolates collected in Quebec.Red and Green masks across subpanel A, indicate loci of possible purifying selection and selective sweeps, respectively.

In Table 2b, we show the average estimate of the nucleotide diversity and Watterson’s theta on the basis of 5 kb windows together with the test statistic for Tajima’s D and Fay and Wu’s H. The local estimates of these test statistics across the genome can be found in Figure 4. Interestingly we also show a much higher estimate of Fay and Wu’s H for the European population (−1.04) compared to the populations sampled from the Americas (−0.06,-0.47). This means that the European population has an excess of high-frequency derived SNPs (with *N. major* as ancestral species) which can be caused by selective sweeps (Sterken *et al*., 2009), but selection works locally, whereas the demographic history affects the whole genome (Cavalli-Sforza, 1966). The European population have negative H values across the genome indicating a residual pattern after a bottleneck. Figure 4 shows Fey’s and Wu’s H.

**Table 2(b).** Window statistics incl frequency based neutrality test statistics.

| Origin | n | $\theta_w$ (5kb) | $\theta_\pi$ (5kb) | Tajima's D (5kb) | Fay & Wu's H (5kb) |
| --- | --- | --- | --- | --- | --- |
| Europe | 43 | 3.9654 | 3.6983 | -0.3252 | -1.0478 |
| Quebec | 15 | 2.9009 | 2.6835 | -0.3824 | -0.0620 |
| BC | 6 | 4.9305 | 4.3286 | -0.6323 | -0.468 |

For the three populations we performed window based analyses (5kb) for Watterson's theta, the pairwise theta along with Tajima's D and Fay and Wu's H. Due to the unevenness of coverage for the different windows we discarded those windows that had less than the average number of sites per window, this was done for each chromosome and each population. Again we remark that Fay and Wu's H requires proper polarization to the ancestral states whereas the Tajima's D is invariant with regards to the choice of polarization.

Disregarding the sample size difference, the SFS (Figure S2) of the Quebec and European populations are very similar in spite of the relatively high F_ST_ of 0.68 between them which is likely driven by the number of fixed differences.

The majority of polymorphic sites called with GATK in the Quebec lineage (61%) are not polymorphic in the European lineage. Similarly, 89% of the polymorphic sites in Europe are private for the European population, and thus only observed in Europe.

### Time to the most recent common ancestor

The split between the British Columbian lineage and the other sampled lineages of *N. neomacrospora* was estimated by the Bayesian analysis to have occurred around ten thousand years before present. The time estimate comes with wide confidence intervals, the 95% highest posterior density (HPD) of the estimate includes a split estimate of 96 kyr BP. The Chinese lineage diverged within the last 79 kyr, with an estimated most likely date around eight kyr BP. The two closest related lineages, the European and the Quebec lineages, split into two separate lineages some two thousand years ago. The 95% HPD of this last estimate is from 200 to 20,200 ybp (All HPD values can be found in Figure S6). If the divergence analysis was performed under the assumption of exponential population growth (Coalescent Exponential Population model in Beast), all median divergence times are roughly halved, and the upper 95% HPDs are divided by four. This gives a median divergence time between the European and Quebec lineages of approximately one thousand years ago.

A mutation rate of 2.44 x 10^−7^ nucleotides per year was estimated using BEAST. Based on this mutation rate, and the 2D-SFS, the split time estimated by diffusion approximation with Moments, under the assumption of constant population size, is 22 kyr ago. This estimate falls outside the 95% HPDs of the Bayesian estimate, and pushes the population split further back in time.

### Demographic history

The demographic history was estimated from the joint site frequency spectrum of the Europe and Quebec populations. The four models tested, ranked as follows: no growth, growth only in QC, growth only in EU, and growth in both populations, with the following likelihoods - 19081, −18355, −14746, −11140, respectively. The model allowing for growth in both populations fitted data best and is shown in Figure 5 (see Figure S8, for details on all models). In all four models, we find population size in Quebec higher than in Europe. Further, the estimated demography suggests that the migration after the population split was highly skewed, with the direction of migration predominately going from the Quebec to the European population. The migration is estimated to be four orders of magnitude higher, with 0.391 compared to 3.8 x 10^−5^ events per generation.

**Figure 5.**
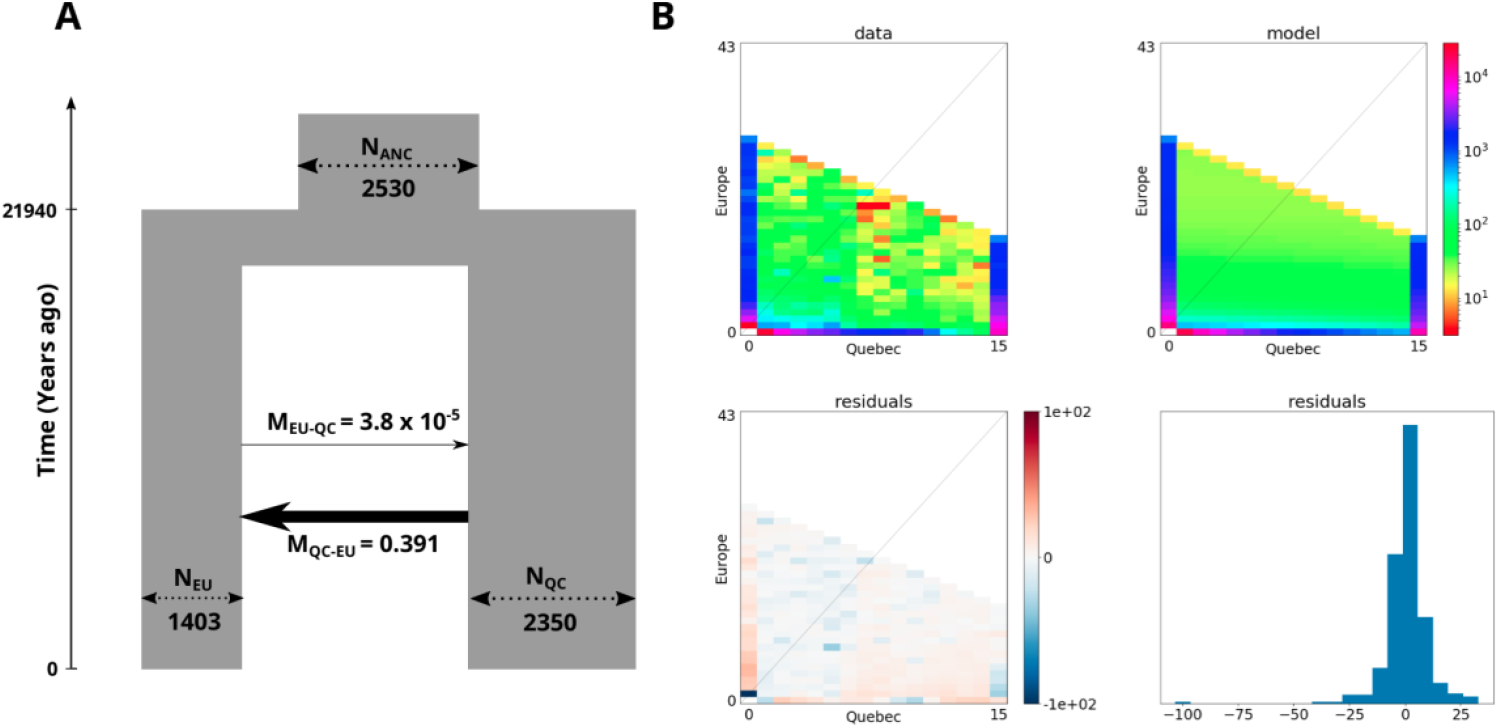
Estimated demography of Europe and Quebec populations. a) Width of boxes represents effective populations sizes and thickness of the black arrows represent the migration rates. b) Data consists of the folded joint (2D) site frequency spectrum of the Quebec and European sample of *Neonectria neomacrospora*. The model fit is given to the right of the SFS, residual of data and model are given below.

Tajima’s D is a SFS based neutrality test statistic sensitive to selection and population size changes. Positive values of Tajima’s D are interpreted to indicate balancing selection and/or decreasing population size, values near zero indicate neutrality, and negative values indicate an excess of rare alleles resulting from a selective sweep, recent population expansion or purifying selection (Tajima, 1989). Small sample sizes are, by sampling error, prone to have proportionally fewer rare alleles then the population sampled. This introduces a bias in Watterson’s theta, which carries over to Tajima’s D. Small sample sizes leads to underestimation by Waterson theta, and subsequently, an overestimation of Tajima’s D. We calculated Tajima’s D for the European and Quebec populations and estimated the effect of the different sample sizes by subsampling the larger European sample down to the size of the Quebec sample (n=15). Mean values of Tajima’s D were calculated based on 100 subsamples without replacement (Figure S6). Figure S6a shows that Quebec values primarily falls between −1 and 2 centered slightly to the positive side of zero. Tajima’s D for the European population has a broader distribution including values above 2, in the original sample of N=43 and at n=15. Interestingly, the subsampling of the European sample reshaped the density distribution of Tajima’s D values, rendering a slightly negative peak and a substantially higher proportion of SNPs with a lower frequency than observed in Quebec.

The Bayesian inference of ancestral population sizes, illustrated with the Bayesian skyline plot in figure S7, does not find a significant difference in ancestral median effective population sizes between Quebec and Europe. Only a minor signal of expansion was detected in Quebec, but the European population is estimated by the EBSP analysis to have expanded its effective population size one order over the last 60-80 generations.

### Linkage Disequilibrium

The analysis of linkage disequilibrium (LD) decay across the genome revealed that the pairwise LD in the Quebec population appears to plateau much sooner than the LD in the European population. The mean r^2^ values of the Quebec samples reaches a plateau within 3 kb (r^2^=0.29), the European population in comparison shows markedly higher r^2^ values, and a slope extending beyond 10 kb (Figure 6). Sample size can bias a LD decay analysis resulting in a false bottleneck signal (Rogers, 2014). Thus, we chose ten random subsamples to n=15 of the European sample to mimic the sample size in Quebec. Four out of ten subsamples raised the degree of LD significantly with a delta r^2^ of approximately 0.17 measured at 2-4 kb distance. Below 2 kb distance, the differences diminish; above 4 kb the uncertainty of the estimate increases. Thus, the differences in the rates of LD decay cannot be attributed to different sample sizes in Quebec and Europe. The slow LD decay and the higher amount of LD observed in the European sample are consistent with the presence of a population size bottleneck in the European population.

**Figure 6.**
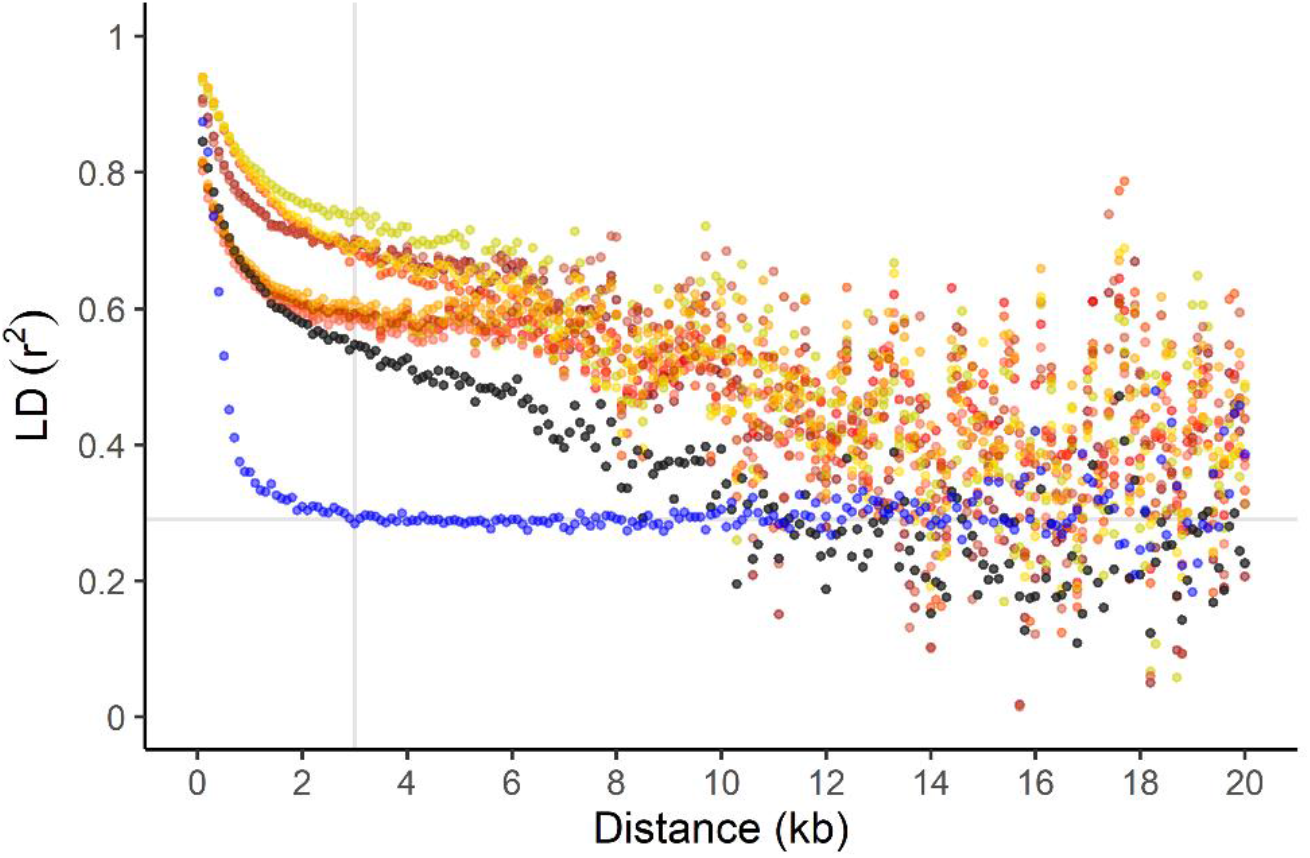
Mean pairwise linkage disequilibrium (r^2^) between polymorphic sites across the genome by distance. Data are stratified by sampling region. Blue: Quebec, n=15. Black: Europe, n=43. Ten independent and random subsamples of the European sample to n= 15 were performed. Nuances of yellow to red are used for the ten subsample. Subsampling were performed to show the effect of sample size, and facilitate a more direct comparison of LD in the Quebec and the European populations.

## DISCUSSION

If the current European epidemic of *N. neomacrospora* had been caused by a recent introduction of the more virulent Quebec lineage of the fungus into Europe, we would then expect that this lineage had either replaced, or created hybrids, that were distinct from pre-epidemic European strains. Using the samples collected in this study, it has not been possible to delineate pre- and post-epidemic strains, and all European strains seem to share a common evolutionary history. The initial introduction of *N. neomacrospora* to either Quebec or Europe must have been sometime before 1957, the collection year of the oldest European strain sequenced in this study. Thus, even though we found that the European and Quebec lineages are phylogenetically closely related in comparison to the strains from British Columbia and China, we cannot support the hypothesis that the current European epidemic is caused by an introduction from Quebec to Europe in the time since the described outbreak in Quebec.

While we do not see evidence of any recent migration, substantial migration from the Quebec population to European population was detected. Whether this was driven alternatively by trans-Atlantic migration, versus migration between sympatric populations followed by trans-Atlantic immigration, cannot be answered using the available data. The migration could have been the result of anthropogenic long-distance dispersal via the global trade of plants and seeds. Seeds of *Abies* spp. are imported to Northern Europe, predominantely from around the Black Sea and North America, and seeds have been shown, at least in one case, to carry *Neonectria* (Talgø *et al*., 2010). Possible routes for a natural long-distance, trans-Atlantic dispersal, of fungi that could be considered is driftwood and wind (Golan and Pringle, 2018).

### Clock rate

We used BEAST to estimate a mutation rate of 2.44 x 10^−7^ per year. Filamentous fungi accumulate mutations through continuous mitotic division in the apical space of the advancing mycelium, and this should be noted in the evaluation of reasonable molecular clock rates for phylogenetic studies in filamentous fungi. Ruiz-Roldán et al. (2010) report a mean time of 92 min between nuclear divisions in the hyphal growth of *Fusarium oxysporium*. This study investigates the germination face and mentions that the rate of nuclear division slows with time. 92 min per cycle equals 5700 mitotic cycles per year. If the true number is between 1000 and 4000 division per year, and the dynamics can be transferred to *N. neomacrospora*, then this approximately equals a mutation rate of 2.5 x 10^−10^ to 1 x 10^−11^ per site per mitosis. Based on the genome sequencing of multiple mutation accumulation lines of *Aspergillus* (Álvarez-Escribano *et al*., 2019) estimated the mutation rate to be 1.1 x 10^−11^ per site per mitosis in *A. fumigatus and* 4.2 x 10^−11^ per site per mitosis in *A. flavus*. Mutations were allowed to accumulate across ∼4000 mitoses (in 30 weeks). Nuclear division rates are influenced by nutrient availability (Ruiz-Roldán *et al*., 2010), and it is difficult to extrapolate from laboratory experiments to field dynamics, but in the light of the above, the mutation rate calculated based on the sampling dates of historical and contemporary samples within this study seem credible.

### Linkage Disequilibrium

The likelihood of recombination between two sites on a chromosome increases with distance, this positive correlation between distance and recombination rate, translate into low recombination rates in the left-hand side of the LD curve, and high in the right-hand. If recombination is the dominating force shaping the genome, and recombination rates are uniform across the genome, then LD blocks will be small and transient.

LD can arise locally as an effect of selection, but genome-wide LD is a result of demographic processes, such as population structure/subdivision, migration and changes in population size (Slatkin, 2008). Population contractions will, in general, lead to the loss of rare haplotypes and raising the genome-wide LD. Genome-wide high LD in one population compared to another have been used to indicate a past bottleneck (Zhang *et al*., 2004).

The steep LD decay and short haplotype blocks observed in Quebec is consistent with a large recombining population. The plateau observed in the same population is proposed, to some extent, to be the background LD caused by somatic mutations. The partial clonal propagation through conidia decreases the effective population size, leading to elevated drift. Drift, although it is stochastic, cause LD uniformly across the genome (Rogers, 2014), since it is not just single alleles, but complete strains, that are lost for future generations. Finally, the background LD can be an effect of the structure detected in the admixture analyses.

The non-random association of SNPs in the European population is an effect of demographic processes since all 10 kb windows analysed across the genome show the same pattern. Since no population structure was detected within Europe, we concentrate on the other possible explanations. We have mentioned population contraction and migration as possible explanations for the observed LD pattern. The negligible effect of drift during a population expanding should, according to Rogers (2014) produce a similar LD curve, and could also contribute to the LD pattern.

When we refer to population contractions, bottlenecks or founder effects, it is often as synonyms for a reduction in effective population size. However, if a few individuals through gained fitness start a population expansion and replace the old diverse population, then we should see a reduction in effective population size, high LD, and an excess of rare alleles not purged by drift.

The high LD in Europe is consistent with positive Tajima’s D values observed. Tajima’s D becomes progressively positive as variation is concentrated on a relatively lower number of segregating sites. Small sample sizes will affect the resolutions of the SFS by underrepresenting rare alleles. This effect is most pronounced in populations in exponential growth or in genes under purifying selection that is characterized by an excess of rare alleles. While nucleotide diversity π is unaffected by sample size, Subramanian (2016) showed via simulation that exponential growth, contrary to constant growth, introduces a bias that renders Waterson θ positively correlated with sample size, with a derived negative correlation between sample size and Tajima’s D. This means that if the population is in exponential growth then the Tajima’s D statistics of the larger sample size (n=43) should be negatively screwed compared to the Tajima’s D of the subsampled population (n=15). What we observed subsampling the European population was a decreased variance, with a reduction of both positive and negative extreme values (Figure S6), and a lowered mean as expected.

The demographic analysis found that the Quebec population originates from an ancestral population larger than the ancestral population that could be inferred from the European population. The smaller ancestral population inferred from the European sample can be caused by a severe bottleneck purging the population of variation present in an ancestral European population. We have shown that the Quebec and European population have a common history expressed in the monophyletic clade of the two populations in the species. It is possible that the two populations diverged sympatrically, or that the split was formed by multiple minor migrations to European leaving a signature of genetic drift.

### Population growth

SFS-based and sequence-based methods have different strengths and weaknesses for demographic inference, some of which comes down to the differences in assumptions and complexity of the models analysed (Schraiber and Akey, 2015; Beichman, Phung and Lohmueller, 2017). Sequence based methods that infer population sizes and demographic events by estimating the rates of coalescence across the genome are insensitive to recent demographic events. In particular, recent demographic events that occurred within the last ∼500-1000 generations have not had enough time to leave their imprint on the genomes in terms of coalescence events. In contrast, SFS based methods are robust to recent changes in demography and can be used to reconstruct both recent and old demographic events. Nevertheless there are some shortcomings to SFS based methods, viz., i) one needs high sample sizes and abundant data to estimate the SFS accurately, and ii) the demographic parameters estimated are constrained by the family of models specified *a priori*.

In this study, we estimated the demography of the European and Quebec samples under four different demographic models, with and without population expansion in the two populations after their split. In all four models, we find higher population sizes in Quebec and a biased migration from Quebec to Europe, suggesting the robustness of these findings to model misspecification. Further, the models that allow for growth in either the European or both populations fits substantially better than the model that does not allow any population growth. Considering these results in combination with the results from LD decay and neutrality statistics, strongly suggests that the European population underwent a population expansion, mostly likely preceded by a founding event.

The Tajima’s D values calculated in windows across the genome show a higher variance when calculated for the European population than it does for the Quebec population. A difference that persists when we look at random subsamples of the European sample. Parts of the European genomes have high D values as described above, but a larger proportion has negative values (Figure 4), indicative of a population expansion. Similarly, did the Extended Bayesian Skyline Plot coalescence analysis estimate a three order of magnitude increase in effective population size within the last 60 years within the European population. These results further support the conclusion that the European population underwent a recent expansion.

We have in this study inferred parts of the demographic history of *N. neomacrospora* and the genetic history of the current European outbreak. When the damage caused by *N. neomacrospora* in Quebec was reported in 1966 (Ouellette and Bard, 1966), and investigations into the cause started, the depth of cankers showed that the initial infection had started at least 28 years earlier. It is reasonable to think that the current epidemic of *N. neomacrospora* in Europe started well before anyone noticed it. We have seen severe damage for at least a decade now; if we to that add the three decades it took to notice the outbreak in Quebec, then we are not far from the 60 years of population growth estimated in this study. The growth within the European population is an important finding. Although seemingly trivial, with an increasing number of reports, in an increasing number of countries, confirming that the population is expanding simplifies the story. An increase in damage caused by *N. neomacrospora* could alternatively have been driven by factors such as climate change, or a increased rate of coinfection by other organisms, altering the interactions between the hosts and a constant fungal population. It is still possible that external factors interact with *N. neomacrospora* to cause the epidemic, but we can conclude that it is at least in part caused by the spread of the fungus.

This study is the first of its kind on *N. neomacrospora*, and was, as such, planned without prior knowledge of the genetic relationship between the geographic populations. Future research should broaden the geographic sampling and identify new populations and borders to the known ones.

## URLS

Beast2, https://www.beast2.org/; FigTree, http://tree.bio.ed.ac.uk/software/figtree/; Funannotate pipeline, https://funannotate.readthedocs.io/en/latest/index.html; PLINK, https://www.cog-genomics.org/plink/1.9/

## ACKNOWLEDGEMENT

We thank Dr Wen-Ying Zhuang (Chinese Academy of Sciences, Beijing) for providing an isolate of *N. neomacrospora* from China. Anne Uimari (Natural Resources Institute Finland, Luke) for collecting and providing samples from Finland. Halvor Solheim, Venche Talgø and Jan-Ole Skage for samples from Norway. Sophie Schmitz (Walloon Agricultural Research Centre) provided an isolate from Belgium.

We thank the Danish National High-Throughput DNA Sequencing Centre for its services. The Danish Christmas Tree Association supported fieldwork and sequencing that made this work possible.

## DATA AVAILABILITY STATEMENT

Raw reads and genomes assemblies of the 71 isolates described in this study are available the European Nucleotide Archive under the study accession number: PRJEB41540. The authors declare that all data of this study are available from the corresponding author upon reasonable request.

## Notes

### Competing Interest Statement

The authors have declared no competing interest.

