## Supplemental material for "Population genomics of the emerging forest pathogen *Neonectria neomacrospora*"

### Supplemental figures

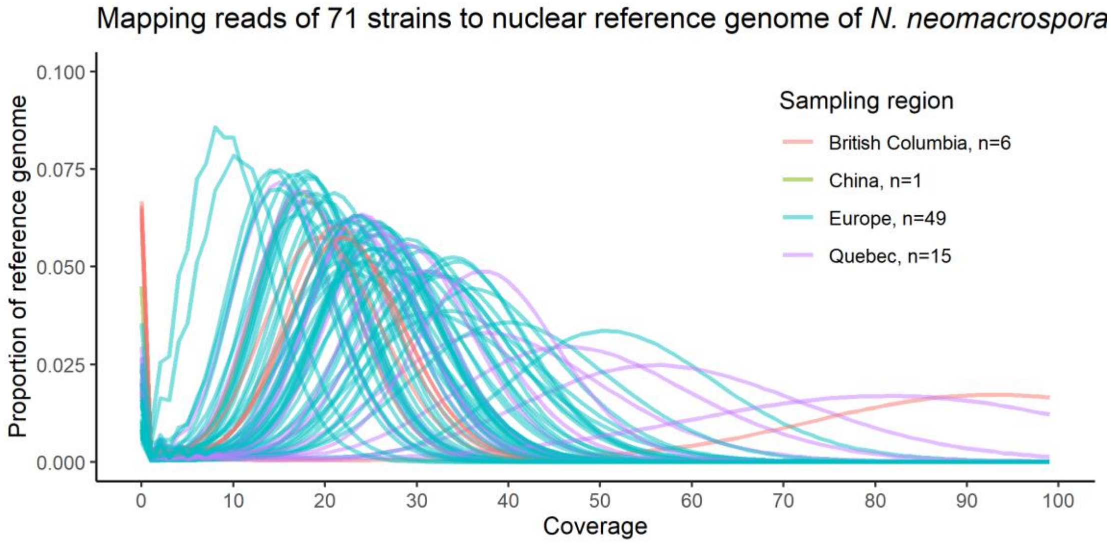


**Figure S1 | Sequencing depth of single isolate mapped to reference genome.** 150 PE Illumina shotgun reads for each stain was map to the gapless genome of Neonectria neomacrospora strain KNNDK1. Coverage was calculated per nucleotide of reference, and proportion of the reference genome was summarised as a function of sequencing depth (coverage). Coverage was calculated using the genomecov option with bedtools and visualized in R. No hump or shoulders can be seen on the right side of any of the 711 curves. Thus, no evidence of chromosomal aneuploidy was found.

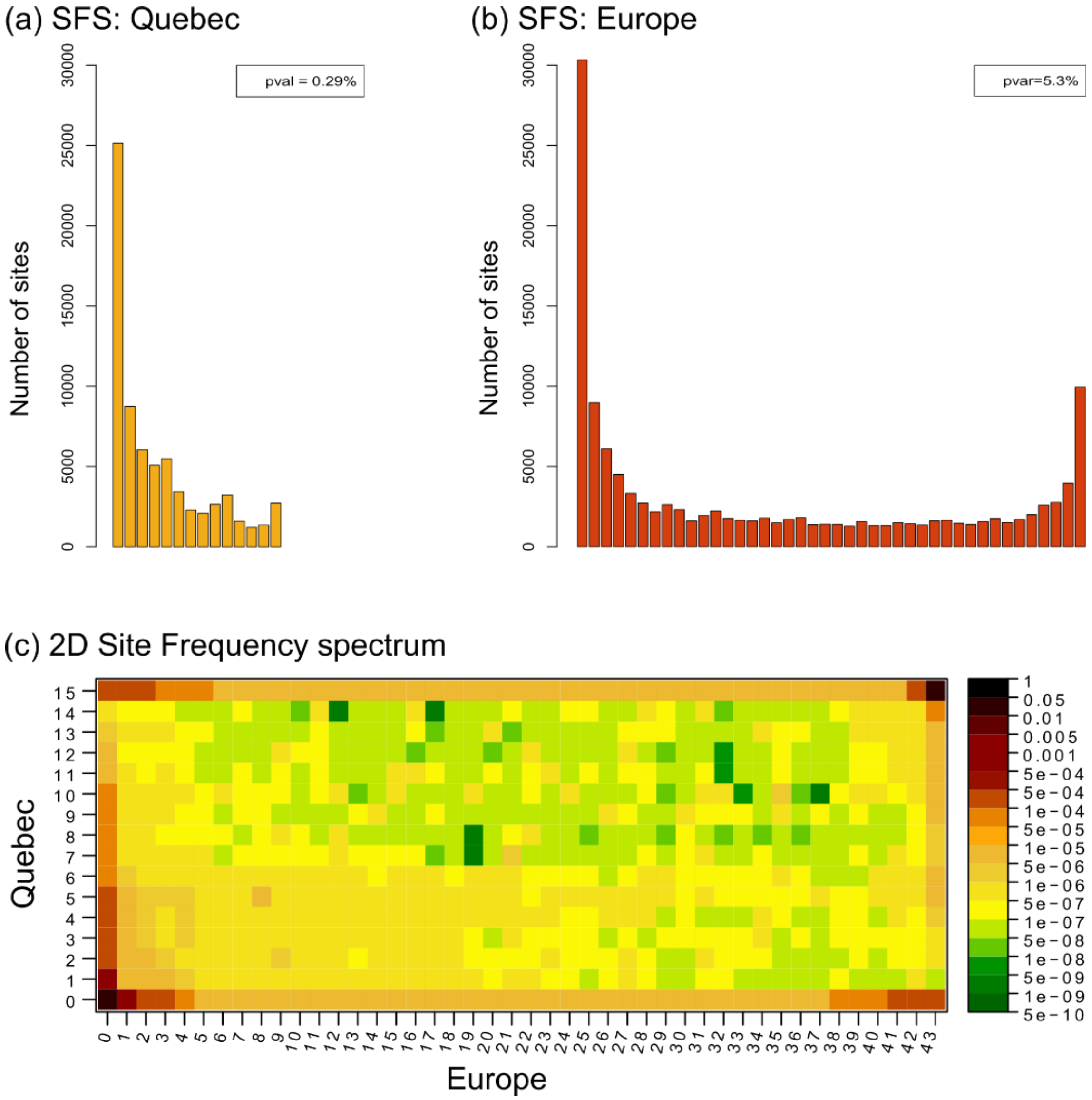


**Figure S2 | Site Frequency Spectrums (SFS) for *Neonectria neomacrospora*.** a) and b), SFS's for the Quebec and European population samples. c) The 2D (joint) SFS of the Quebec and European populations.

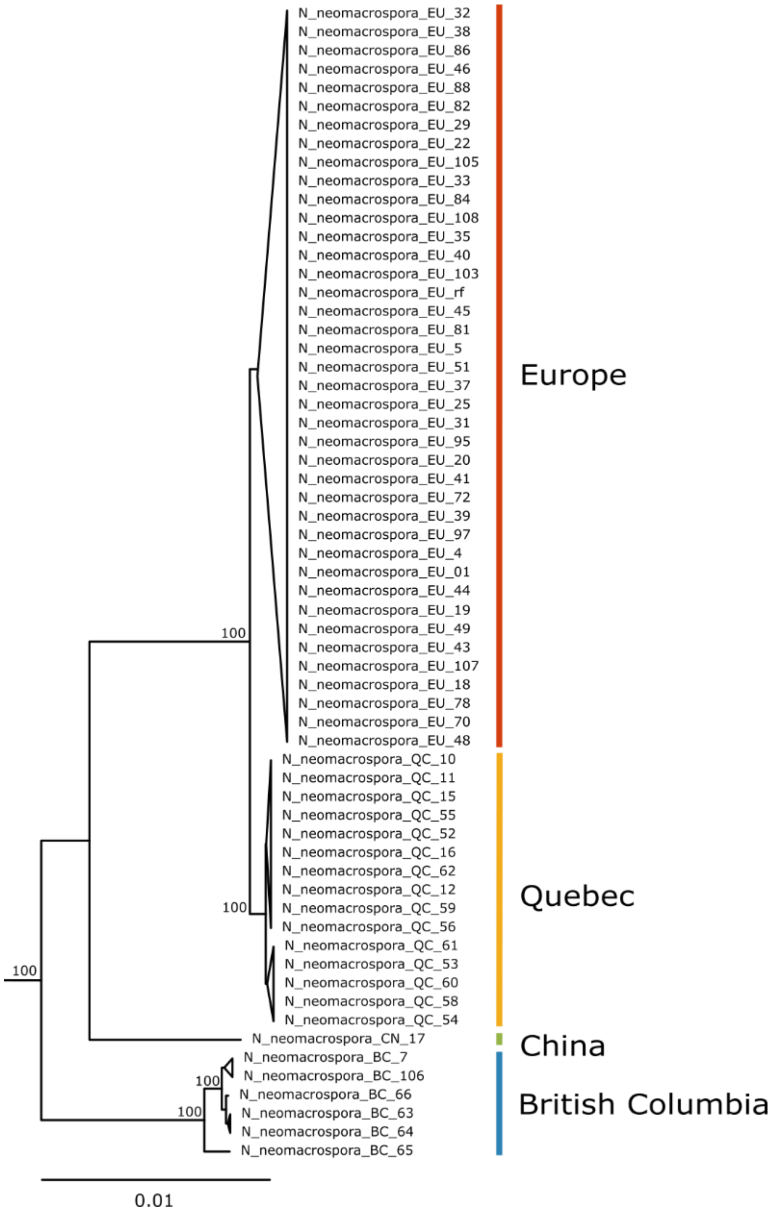


**Figure S3 | Maximum-likelihood phylogeny.** The phylogeny is based on 51 single-copy genes dispersed across the genome with a minimum distance of 10 kb. Bootstrap support form 100 trees are given at nodes.

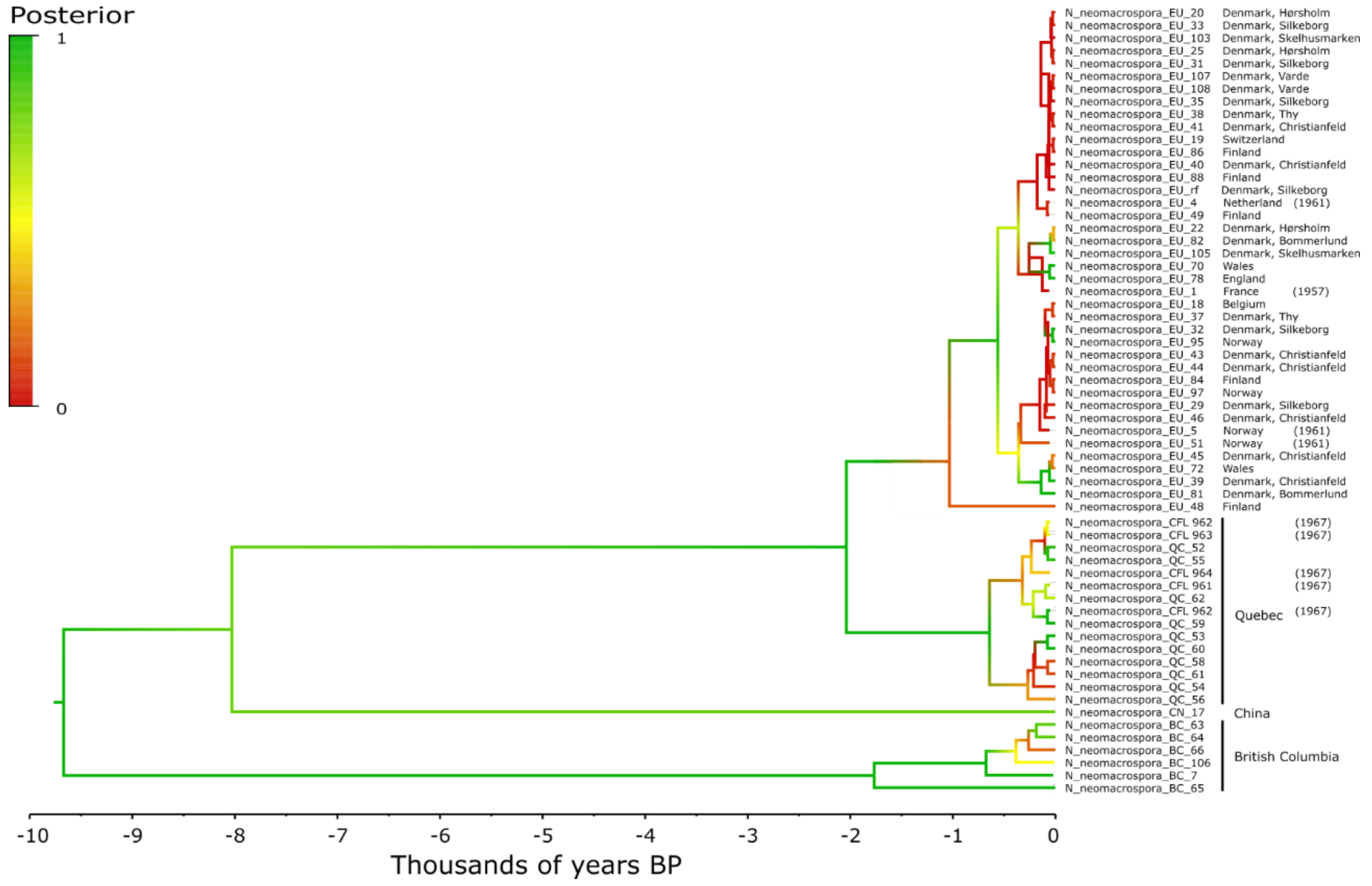


**Figure S4 | Phylogenetic tree** of 62 *Neonectria neomacrospora* strains from Europe (n=40), Quebec (n=15), China (n=1) and British Columbia (n=6).. Branch colours indicate posterior values.

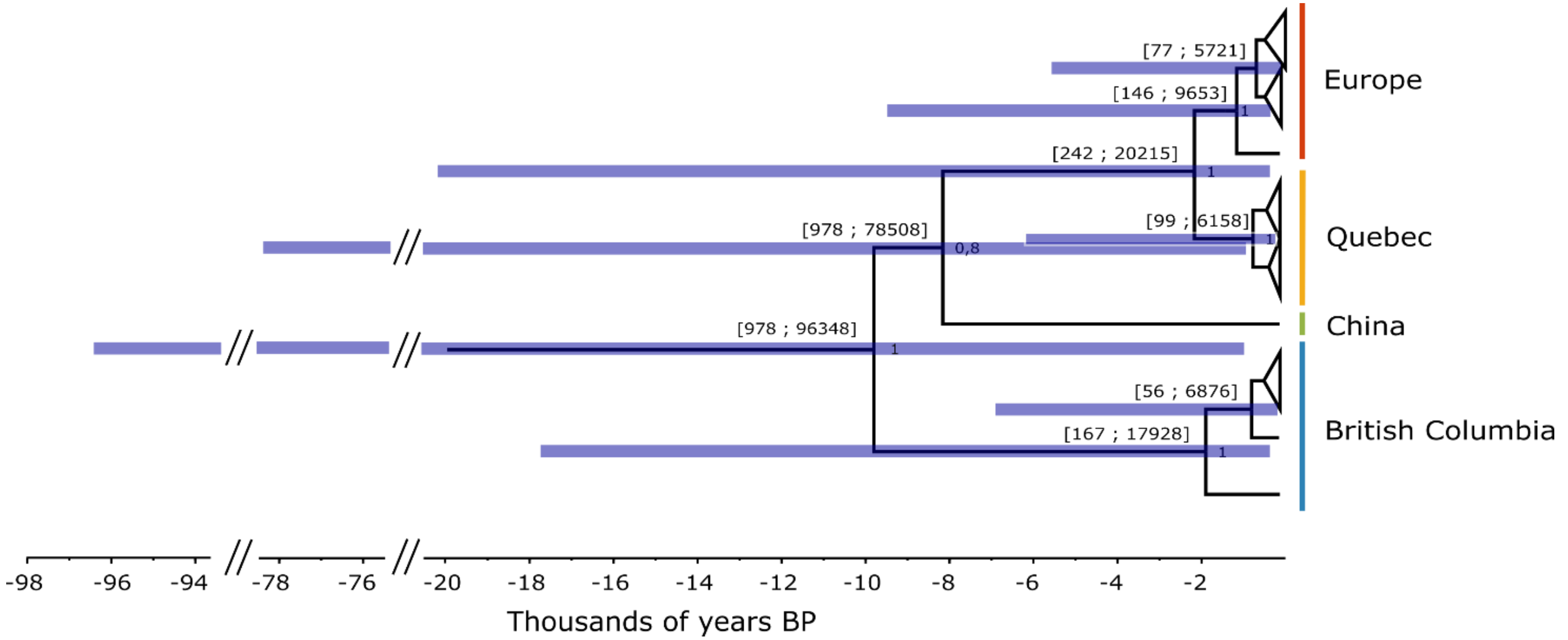


**Figure S5 | Phylogenetic tree** of 62 *Neonectria neomacrospora* strains from Europe (n=40), Quebec (n=15), China (n=1) and British Columbia (n=6). Branch colours indicate posterior values. Blue bars and numbers in brackes indicate the 95% HPD of the node ages.

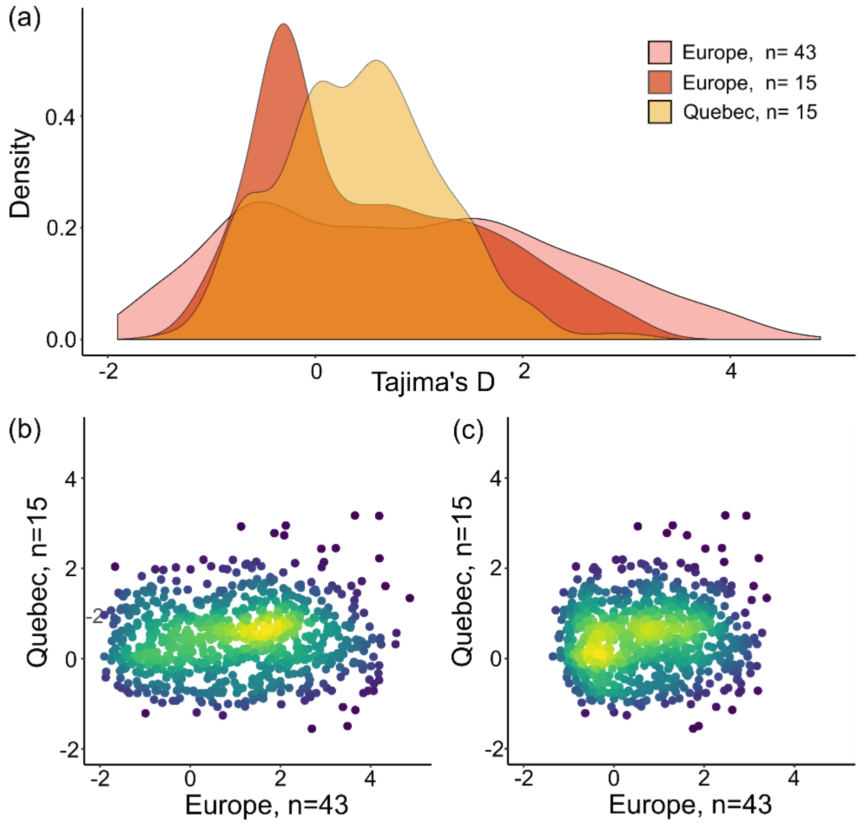


**Figure S6 | Tajima's D for the Quebec and European populations of *N. neomacrospora*.** The European population was subsampled to match the Quebec sample size of N=15. Tajima's D was calculated in windows of 10 kb across the genome, 824 windows had SNPs in both population. Rarefaction was done by random subsampling of 15 strains, the mean values reported are based on 100 rarefactions. A) The density distribution of Tajima's D values for the three stratifications, 1. complete European sample clone-corrected (1957-2019, n=43), 2. the latter subsampled to n=15, and finally 3. Quebec (1967-2018, n=15). B and C) biplot of Tajima's D for identical genomic windows between isolates.

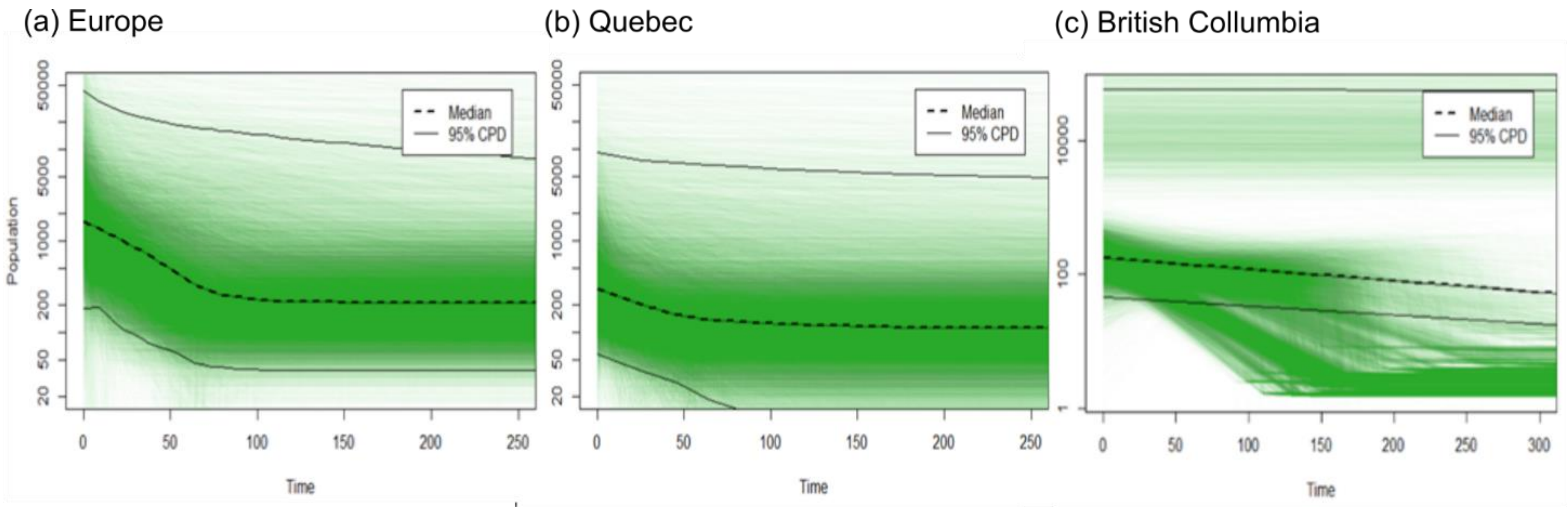


**Figure S7 | Extended Bayeasian Skyline Plots (EBSP).** The EBSP show an increase in effective population size in the European population of *Neonectria neomacrospora*. The x-axis is time measured in generations, and y-axis is equal to $N_e\tau$ (the product of the effective population size and the generation length). The dashed line is the median estimate, where solid lines mark the 95% central posterior density (CPD).

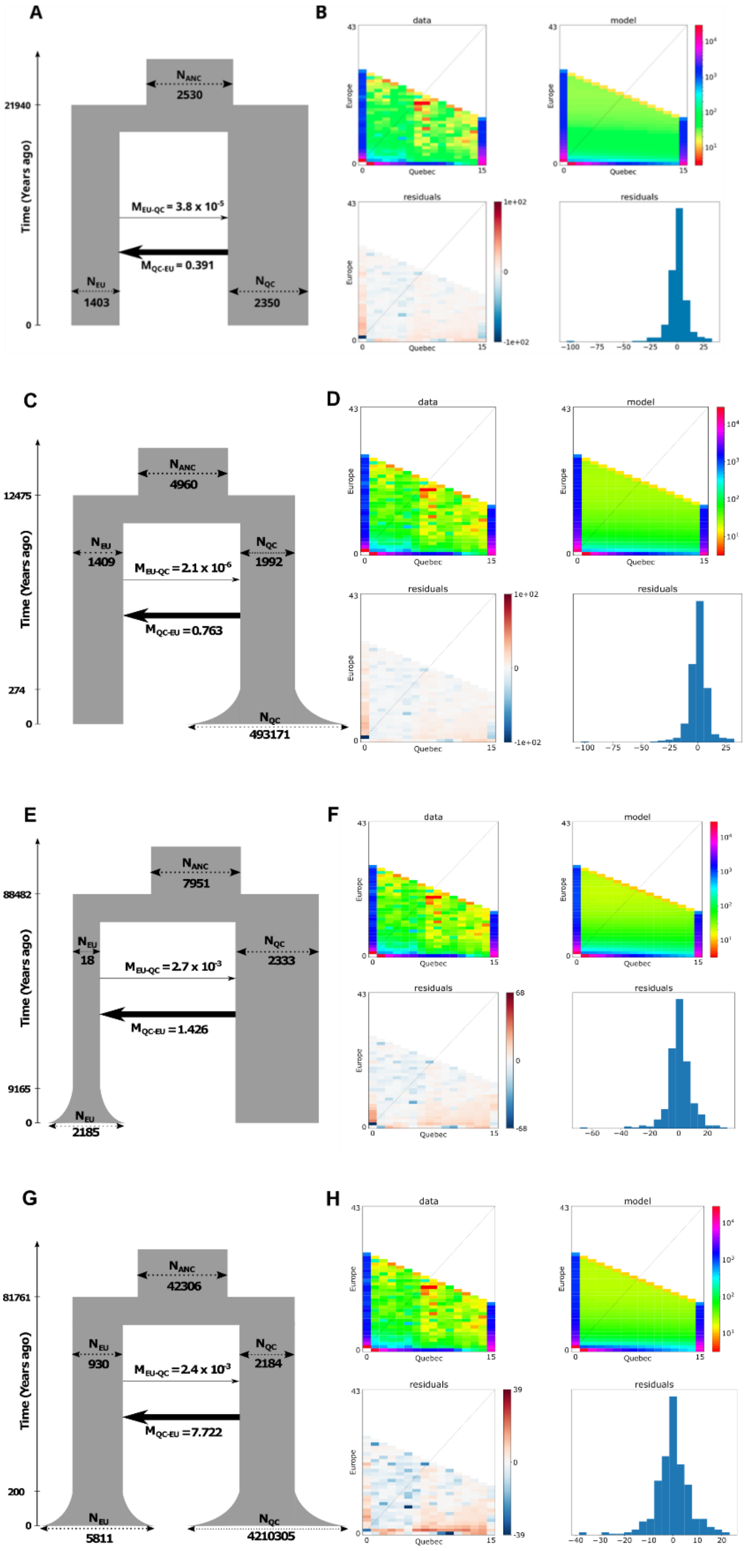


**Figure S8 | Estimated demography of Europe and Quebec populations.** Four models, all an asymmetric migration: a-b) growth in both populations, c-d) growth only in QC, e-f) growth only in EU, or g-h) a constant population size in both populations (i.e. no growth). Scematic models (a,c,e,g): Width of boxes represents effective populations sizes and thickness of the black arrows represent the migration rates. Data-plot (b,d,f,h) Data consists of the folded joint (2D) site frequency spectrum of the Quebec and European sample of *Neonectria neomacrospora*. The model fit is given to the right of the SFS, residual of data and model are given below.

### Table S1 | Assembly statistics and read coverage

| Species | Country | Province | Collected | Source / Assesion no. | ID | Coverage | #contigs | length | GC | N50 | N75 | L50 |
|---|---|---|---|---|---|---|---|---|---|---|---|---|
| *Neonectria neomacrospora* | France | Lac de Longmer, | 1957 | CBS 189.61 | 001 | 54.9 | 793 | 37021198 | 53.64 | 225535 | 135552 | 48 |
| *Neonectria neomacrospora* | Netherlands | Zwolle | 1961 | CBS 324.61 | 004 | 25.7 | 1015 | 37091102 | 53.65 | 236725 | 119474 | 53 |
| *Neonectria neomacrospora* | Belgium | Herbeumont | 2017 | BE5104 | 018 | 20.2 | 1917 | 36461970 | 54.28 | 65786 | 35561 | 170 |
| *Neonectria neomacrospora* | Switzerland | Jura | 2017 | CH01011 | 019 | 20.6 | 895 | 37151449 | 53.51 | 175620 | 90111 | 65 |
| *Neonectria neomacrospora* | Norway | Hordaland | 1958 | NO 1883/5 | 049 | 25.1 | 2047 | 36312304 | 54.41 | 54153 | 29584 | 197 |
| *Neonectria neomacrospora* | Norway | Hordaland | 1961 | CBS 503.67 | 005 | 27.0 | 887 | 37185101 | 53.59 | 199846 | 100159 | 56 |
| *Neonectria neomacrospora* | Norway | Hordaland | 1961 | NO 61-62/1 | 051 | 27.1 | 1588 | 36649618 | 54.23 | 131008 | 68412 | 90 |
| *Neonectria neomacrospora* | Norway | Hordaland | 2019 | NO 252125 | 093 | 25.0 | 1448 | 36953685 | 53.93 | 160193 | 85517 | 70 |
| *Neonectria neomacrospora* | Norway | Hordaland | 2019 | NO 252130 | 095 | 27.1 | 1693 | 36615896 | 54.21 | 116187 | 59460 | 95 |
| *Neonectria neomacrospora* | Norway | Hordaland | 2019 | NO 252140 | 097 | 24.0 | 1701 | 36571452 | 54.15 | 130597 | 67827 | 87 |
| *Neonectria neomacrospora* | Denmark | East Zealand | 2015 | DK01011 | 020 | 34.5 | 1223 | 36975782 | 53.79 | 192378 | 90265 | 64 |
| *Neonectria neomacrospora* | Denmark | East Zealand | 2016 | DK01081 | 022 | 19.4 | 1220 | 37053232 | 53.77 | 205333 | 101824 | 58 |
| *Neonectria neomacrospora* | Denmark | East Zealand | 2015 | DK01132 | 025 | 22.0 | 8714 | 38228153 | 53.77 | 117120 | 57303 | 93 |
| *Neonectria neomacrospora* | Denmark | East Jutland | 2016 | DK02073 | 029 | 38.5 | 663 | 37142615 | 53.56 | 291696 | 156980 | 42 |
| *Neonectria neomacrospora* | Denmark | East Jutland | 2016 | DK02232 | 031 | 31.1 | 1094 | 36934545 | 53.74 | 208103 | 117157 | 55 |
| *Neonectria neomacrospora* | Denmark | East Jutland | 2016 | DK02251 | 032 | 31.2 | 926 | 37076846 | 53.66 | 240590 | 135011 | 45 |
| *Neonectria neomacrospora* | Denmark | East Jutland | 2016 | DK02261 | 033 | 36.1 | 1854 | 37325201 | 53.61 | 252208 | 146648 | 47 |
| *Neonectria neomacrospora* | Denmark | East Jutland | 2016 | DK02281 | 035 | 29.0 | 1403 | 36943622 | 53.83 | 211433 | 128047 | 50 |
| *Neonectria neomacrospora* | Denmark | North Jutland | 2015 | DK03011 | 037 | 27.3 | 1552 | 36979382 | 53.82 | 177687 | 93370 | 62 |
| *Neonectria neomacrospora* | Denmark | North Jutland | 2015 | DK03021 | 038 | 27.4 | 1102 | 37173688 | 53.59 | 240501 | 120041 | 46 |
| *Neonectria neomacrospora* | Denmark | Southern Jutland | 2018 | DK10021 | 039 | 27.9 | 918 | 37095600 | 53.53 | 242851 | 139445 | 47 |
| *Neonectria neomacrospora* | Denmark | Southern Jutland | 2018 | DK10041 | 040 | 35.9 | 705 | 37114797 | 53.58 | 298831 | 152924 | 39 |
| *Neonectria neomacrospora* | Denmark | Southern Jutland | 2018 | DK10051 | 041 | 31.0 | 680 | 37269791 | 53.41 | 289122 | 161549 | 41 |
| *Neonectria neomacrospora* | Denmark | Southern Jutland | 2018 | DK10091 | 043 | 19.9 | 1038 | 37075113 | 53.61 | 201219 | 104527 | 53 |
| *Neonectria neomacrospora* | Denmark | Southern Jutland | 2018 | DK10101 | 044 | 27.6 | 1029 | 37100765 | 53.59 | 235967 | 137390 | 48 |
| *Neonectria neomacrospora* | Denmark | Southern Jutland | 2018 | DK10111 | 045 | 25.3 | 713 | 37052615 | 53.59 | 225935 | 122065 | 48 |
| *Neonectria neomacrospora* | Denmark | Southern Jutland | 2018 | DK10121 | 046 | 22.5 | 977 | 37146474 | 53.56 | 216117 | 120842 | 56 |
| *Neonectria neomacrospora* | Denmark | Southern Jutland | 2018 | DK09011 | 081 | 16.5 | 2603 | 36488173 | 54.25 | 45725 | 24190 | 248 |
| *Neonectria neomacrospora* | Denmark | Southern Jutland | 2018 | DK09111 | 082 | 12.4 | 7523 | 36022668 | 54.44 | 8694 | 5077 | 1277 |
| *Neonectria neomacrospora* | Denmark | East Jutland | 2015 | DK04021 | 103 | 35.4 | 701 | 37160680 | 53.50 | 268646 | 164683 | 46 |
| *Neonectria neomacrospora* | Denmark | East Jutland | 2015 | DK04032 | 104 | 34.2 | 1013 | 37161798 | 53.66 | 263382 | 142176 | 45 |
| *Neonectria neomacrospora* | Denmark | East Jutland | 2015 | DK04071 | 105 | 32.7 | 978 | 37110878 | 53.58 | 225360 | 115094 | 51 |
| *Neonectria neomacrospora* | Denmark | West Jutland | 2016 | DK07033 | 107 | 19.8 | 1794 | 36488527 | 54.25 | 106266 | 56441 | 108 |
| *Neonectria neomacrospora* | Denmark | West Jutland | 2016 | DK07041 | 108 | 35.4 | 541 | 37062301 | 53.60 | 261055 | 142735 | 47 |

| | | | | | | | | | | | | |
|---|---|---|---|---|---|---|---|---|---|---|---|---|
| *Neonectria neomacrospora* | Finland | Southern Finland | 2018 | FI01011 | 048 | 19.5 | 1075 | 37203405 | 53.58 | 220690 | 115344 | 52 |
| *Neonectria neomacrospora* | Finland | Southern Finland | 2019 | FI01021 | 084 | 28.7 | 1762 | 36452860 | 54.34 | 78007 | 43594 | 146 |
| *Neonectria neomacrospora* | Finland | Southern Finland | 2019 | FI01041 | 086 | 26.2 | 1841 | 36464338 | 54.26 | 78890 | 43080 | 137 |
| *Neonectria neomacrospora* | Finland | Southern Finland | 2019 | FI01061 | 088 | 21.5 | 998 | 37166452 | 53.57 | 184820 | 94892 | 61 |
| *Neonectria neomacrospora* | Canada | Quebec | 1967 | QFB19253 / | 010 | 60.6 | 1185 | 36695899 | 53.76 | 191600 | 93673 | 56 |
| *Neonectria neomacrospora* | Canada | Quebec | 1967 | QFB19255 / | 011 | 41.3 | 1416 | 36870701 | 53.78 | 143783 | 80213 | 74 |
| *Neonectria neomacrospora* | Canada | Quebec | 1967 | QFB19262 / | 012 | 49.8 | 1391 | 36862386 | 53.71 | 179801 | 90456 | 61 |
| *Neonectria neomacrospora* | Canada | Quebec | 2018 | CA01011 | 052 | 27.7 | 1462 | 36679750 | 53.98 | 149872 | 79672 | 74 |
| *Neonectria neomacrospora* | Canada | Quebec | 2018 | CA01021 | 053 | 25.5 | 1685 | 37291616 | 53.54 | 188252 | 99432 | 55 |
| *Neonectria neomacrospora* | Canada | Quebec | 2018 | CA01031 | 054 | 30.5 | 864 | 37036564 | 53.56 | 223232 | 114296 | 51 |
| *Neonectria neomacrospora* | Canada | Quebec | 2018 | CA01041 | 055 | 83.0 | 972 | 36425792 | 54.20 | 142821 | 84433 | 79 |
| *Neonectria neomacrospora* | Canada | Quebec | 2018 | CA01051 | 056 | 32.7 | 1461 | 36964005 | 53.61 | 98032 | 52487 | 115 |
| *Neonectria neomacrospora* | Canada | Quebec | 2018 | CA01061 | 058 | 39.2 | 869 | 36953958 | 53.61 | 271815 | 132335 | 44 |
| *Neonectria neomacrospora* | Canada | Quebec | 2018 | CA01071 | 059 | 16.4 | 1910 | 36263659 | 54.40 | 58786 | 31512 | 192 |
| *Neonectria neomacrospora* | Canada | Quebec | 2018 | CA01081 | 060 | 24.3 | 985 | 37018946 | 53.61 | 188467 | 106612 | 57 |
| *Neonectria neomacrospora* | Canada | Quebec | 2018 | CA01091 | 061 | 18.7 | 1639 | 36370405 | 54.31 | 97585 | 52049 | 112 |
| *Neonectria neomacrospora* | Canada | Quebec | 1967 | CFL 963 | 015 | 25.5 | 850 | 37098179 | 53.48 | 224687 | 107262 | 56 |
| *Neonectria neomacrospora* | Canada | Quebec | 1967 | CFL 964 | 016 | 20.3 | 1387 | 36969610 | 53.60 | 118973 | 63750 | 93 |
| *Neonectria neomacrospora* | Canada | Quebec | 2018 | CA02011 | 062 | 25.2 | 1605 | 36893067 | 53.82 | 158930 | 87451 | 65 |
| *Neonectria neomacrospora* | Canada | British Columbia | 1996 | CBS 118985 | 007 | 22.3 | 9490 | 42242818 | 50.91 | 76109 | 36528 | 165 |
| *Neonectria neomacrospora* | Canada | British Columbia | 2018 | CA03011 | 063 | 23.1 | 6576 | 39112794 | 52.94 | 64377 | 32101 | 177 |
| *Neonectria neomacrospora* | Canada | British Columbia | 2018 | CA03021 | 064 | 20.8 | 6175 | 38525360 | 53.41 | 50293 | 27196 | 219 |
| *Neonectria neomacrospora* | Canada | British Columbia | 2018 | CA03031 | 065 | 23.0 | 8552 | 40210002 | 52.41 | 56026 | 27875 | 205 |
| *Neonectria neomacrospora* | Canada | British Columbia | 2018 | CA03041 | 066 | 20.4 | 6543 | 39312554 | 52.77 | 61360 | 31090 | 184 |
| *Neonectria neomacrospora* | Canada | British Columbia | 2005 | CBS 118984 | 106 | 89.8 | 2558 | 40869945 | 51.18 | 98013 | 47611 | 125 |
| *Neonectria neomacrospora* | China | Shennongjia | 2014 | HMAS 252906 | 017 | 18.7 | 5484 | 40961935 | 51.39 | 90957 | 48970 | 130 |

**Table S2 | Annotation for the 51 single-copy genes used for phylogenetic inference**

| Chr | Start | End | | Product | Ontology_term | InterProScan and PFAM reference | note |
|---|---|---|---|---|---|---|---|
| chr3 | 4144044 | 4145497 | + | **26S proteasome regulatory subunit 6B** | GO_component: GO:0005737 - cytoplasm [Evidence IEA],GO_function: GO:0016787 - hydrolase activity [Evidence IEA],GO_function: GO:0005524 - ATP binding [Evidence IEA],GO_process: GO:0030163 - protein catabolic process [Evidence IEA] | InterPro:IPR003593,InterPro:IPR003959,InterPro:IPR003960,InterPro:IPR005937,InterPro:IPR027417,InterPro:IPR032501 PFAM:PF00004,PFAM:PF07724,PFAM:PF07728,PFAM:PF16450 | note=EggNog:ENOG410PGYA,COG:O |
| chr2 | 2432671 | 2434064 | - | **26S proteasome regulatory subunit rpn6** | | InterPro:IPR000717,InterPro:IPR036390,InterPro:IPR040773,InterPro:IPR040780,PFAM:PF01399,PFAM:PF18055, PFAM:PF18503 | note=BUSCO:EOG092630YS, EggNog:ENOG410PJ50,COG:O |
| chr2 | 2832680 | 2833181 | - | **40S ribosomal protein S20** | GO_component: GO:0005840 - ribosome [Evidence IEA],GO_component: GO:0015935 - small ribosomal subunit [Evidence IEA],GO_function: GO:0003735 - structural constituent of ribosome [Evidence IEA],GO_function: GO:0003723 - RNA binding [Evidence IEA],GO_process: GO:0006412 - translation [Evidence IEA] | InterPro:IPR001848,InterPro:IPR005729,InterPro:IPR018268,InterPro:IPR027486,InterPro:IPR036838,PFAM:PF00338 | note=BUSCO:EOG09265CFP, EggNog:ENOG410PQ4U,COG:J |
| chr1 | 2534170 | 2534807 | - | **60S ribosomal protein L11** | GO_component: GO:0005840 - ribosome [Evidence IEA],GO_function: GO:0003735 - structural constituent of ribosome [Evidence | InterPro:IPR002132,InterPro:IPR020929,InterPro:IPR022803,InterPro:IPR031309,InterPro:IPR031310 | note=BUSCO:EOG09264X8J, EggNog:ENOG410PGC7,COG:J |

| | | | | | | | |
|---|---|---|---|---|---|---|---|
| | | | | | IEA],GO_process: GO:0006412 - translation [Evidence IEA] | PFAM:PF00281,PFAM:PF00673 | |
| chr2 | 3997720 | 3998500 | - | **60S ribosomal protein L25** | GO_component: GO:0005840 - ribosome [Evidence IEA],GO_function: GO:0003735 - structural constituent of ribosome [Evidence IEA],GO_process: GO:0006412 - translation [Evidence IEA] | InterPro:IPR005633,InterPro:IPR012677,InterPro:IPR012678,InterPro:IPR013025,PFAM:PF00276,PFAM:PF03939 | note=BUSCO:EOG092657YR,EggNog:ENOG410PPAS,COG:J |
| chr2 | 4488547 | 4489250 | - | **60S ribosomal protein L32** | GO_component: GO:0005840 - ribosome [Evidence IEA],GO_function: GO:0003735 - structural constituent of ribosome [Evidence IEA],GO_process: GO:0006412 - translation [Evidence IEA] | InterPro:IPR001515,InterPro:IPR036351,PFAM:PF01655 | note=EggNog:ENOG410PNPG,COG:J |
| chr5 | 2784402 | 2785147 | - | **ATP synthase d subunit** | GO_component: GO:0000276 - mitochondrial proton-transporting ATP synthase complex, coupling factor F(o) [Evidence IEA],GO_function: GO:0015078 - proton transmembrane transporter activity [Evidence IEA],GO_process: GO:0015986 - ATP synthesis coupled proton transport [Evidence IEA] | InterPro:IPR008689,InterPro:IPR036228,PFAM:PF05873 | note=EggNog:ENOG410PN6K,COG:C |
| chr7 | 781789 | 784130 | - | **bifunctional tryptophan synthase trp1** | GO_function: GO:0003824 - catalytic activity [Evidence IEA],GO_function: GO:0004640 - phosphoribosylanthranilate isomerase activity [Evidence IEA],GO_function: GO:0004425 - indole-3-glycerol-phosphate synthase activity [Evidence IEA],GO_function: GO:0004049 - anthranilate synthase activity [Evidence IEA],GO_process: GO:0006568 - tryptophan metabolic process [Evidence IEA] | InterPro:IPR001240,InterPro:IPR001468,InterPro:IPR006221,InterPro:IPR011060,InterPro:IPR013785,InterPro:IPR013798,InterPro:IPR016302, InterPro:IPR017926,InterPro:IPR029062, PFAM:PF00117,PFAM:PF00218,PFAM:PF00697 | note=BUSCO:EOG09261OLD,MEROPS:MER0045094,EggNog:ENOG410PGPM,COG:E |

| | | | | | | | |
|---|---|---|---|---|---|---|---|
| chr3 | 2368442 | 2369399 | + | **diphthine synthase** | GO_function: GO:0004164 - diphthine synthase activity [Evidence IEA],GO_function: GO:0008168 - methyltransferase activity [Evidence IEA],GO_process: GO:0017183 - peptidyl-diphthamide biosynthetic process from peptidyl-histidine [Evidence IEA] | InterPro:IPR000878,InterPro:IPR004551,InterPro:IPR014776,InterPro:IPR014777,InterPro:IPR035996,PFAM:PF00590 | note=BUSCO:EOG09263JW5,EggNog:ENOG410PGKF,COG:J |
| chr3 | 4044603 | 4045909 | - | **Elongation of fatty acids protein 2** | GO_function: GO:0004518 - nuclease activity [Evidence IEA],GO_function: GO:0003677 - DNA binding [Evidence IEA],GO_function: GO:0003824 - catalytic activity [Evidence IEA],GO_function: GO:0016788 - hydrolase activity, acting on ester bonds [Evidence IEA],GO_process: GO:0006281 - DNA repair [Evidence IEA] | InterPro:IPR006084,InterPro:IPR006085,InterPro:IPR006086,InterPro:IPR008918,InterPro:IPR019974,InterPro:IPR023426,InterPro:IPR029060,InterPro:IPR036279,PFAM:PF00752,PFAM:PF00867 | note=BUSCO:EOG092634B1,EggNog:ENOG410PFFR,COG:L |
| chr8 | 662079 | 663125 | + | **Eukaryotic translation initiation factor 6** | GO_function: GO:0043022 - ribosome binding [Evidence IEA],GO_process: GO:0042256 - mature ribosome assembly [Evidence IEA] | InterPro:IPR002769,PFAM:PF01912 | note=BUSCO:EOG092644O2,EggNog:ENOG410PFYZ,COG:J |
| chr1 | 2503148 | 2504115 | + | **GTP-binding nuclear protein gsp1/Ran** | GO_function: GO:0005525 - GTP binding [Evidence IEA],GO_function: GO:0003924 - GTPase activity [Evidence IEA],GO_process: GO:0006913 - nucleocytoplasmic transport [Evidence IEA] | InterPro:IPR001806,InterPro:IPR002041,InterPro:IPR005225,InterPro:IPR027417,PFAM:PF00025,PFAM:PF00071,PFAM:PF08477 | note=BUSCO:EOG09264LKR,EggNog:ENOG410PFRI,COG:U |

| | | | | | | | |
|---|---|---|---|---|---|---|---|
| chr7 | 770316 | 771581 | + | **guanine nucleotide-binding protein subunit alpha** | GO_component: GO:0005834 - heterotrimeric G-protein complex [Evidence IEA],GO_function: GO:0003924 - GTPase activity [Evidence IEA],GO_function: GO:0031683 - G-protein beta/gamma-subunit complex binding [Evidence IEA],GO_function: GO:0019001 - guanyl nucleotide binding [Evidence IEA],GO_function: GO:0005525 - GTP binding [Evidence IEA],GO_function: GO:0001664 - G protein-coupled receptor binding [Evidence IEA],GO_process: GO:0007186 - G protein-coupled receptor signaling pathway [Evidence IEA],GO_process: GO:0007165 - signal transduction [Evidence IEA] | InterPro:IPR001019,InterPro:IPR002975,InterPro:IPR011025,InterPro:IPR027417,PFAM:PF00025,PFAM:PF00503 | note=EggNog:ENOG410PHS2,COG:D,T |
| chr2 | 3857226 | 3857813 | + | **histone H2A** | GO_component: GO:0005634 - nucleus [Evidence IEA],GO_component: GO:0000786 - nucleosome [Evidence IEA],GO_function: GO:0003677 - DNA binding [Evidence IEA],GO_function: GO:0046982 - protein heterodimerization activity [Evidence IEA] | InterPro:IPR002119,InterPro:IPR007125,InterPro:IPR009072,InterPro:IPR032454,InterPro:IPR032458,PFAM:PF00125,PFAM:PF00808,PFAM:PF16211 | note=EggNog:ENOG410PNTK,COG:B |
| chr1 | 1603192 | 1603548 | + | **hypothetical protein** | | | note=EggNog:ENOG410QA9F (Taxomomic profile: 100% Hypocreales, 7 species) |
| chr1 | 2753059 | 2753857 | + | **hypothetical protein** | GO_component: GO:0000139 - Golgi membrane [Evidence IEA],GO_component: GO:0016021 - integral component of membrane [Evidence IEA],GO_component: GO:0005801 - cis-Golgi network [Evidence IEA],GO_process: GO:0006888 | InterPro:IPR023601,PFAM:PF12352 | note=BUSCO:EOG09265AL8,EggNog:ENOG410PM03,COG:U |

| | | | | | | | |
|---|---|---|---|---|---|---|---|
| | | | | | - endoplasmic reticulum to Golgi vesicle-mediated transport [Evidence IEA] | | |
| chr1 | 4079298 | 4080111 | - | **hypothetical protein** | GO_component: GO:0089701 - U2AF [Evidence IEA],GO_function: GO:0003723 - RNA binding [Evidence IEA],GO_function: GO:0046872 - metal ion binding [Evidence IEA],GO_function: GO:0003676 - nucleic acid binding [Evidence IEA],GO_process: GO:0000398 - mRNA splicing, via spliceosome [Evidence IEA] | InterPro:IPR000504,InterPro:IPR000571,InterPro:IPR009145,InterPro:IPR012677,InterPro:IPR035979,PFAM:PF00642 | note=EggNog:ENOG410PHGA,COG:A |
| chr1 | 4414522 | 4415168 | + | **hypothetical protein** | GO_component: GO:0005840 - ribosome [Evidence IEA],GO_function: GO:0003735 - structural constituent of ribosome [Evidence IEA],GO_function: GO:0003723 - RNA binding [Evidence IEA],GO_function: GO:0003676 - nucleic acid binding [Evidence IEA],GO_process: GO:0006412 - translation [Evidence IEA] | InterPro:IPR001892,InterPro:IPR010979,InterPro:IPR018269,InterPro:IPR027437,PFAM:PF00416 | note=BUSCO:EOG09264YEG,EggNog:ENOG410PG18,COG:J |
| chr1 | 5003097 | 5004242 | + | **hypothetical protein** | GO_function: GO:0016787 - hydrolase activity [Evidence IEA] | InterPro:IPR006680,InterPro:IPR032466,PFAM:PF04909 | note=EggNog:ENOG410PH41,COG:S |
| chr1 | 5042192 | 5042930 | - | **hypothetical protein** | GO_component: GO:0016021 - integral component of membrane [Evidence IEA] | InterPro:IPR004932,PFAM:PF03248 | note=BUSCO:EOG09265ANI,EggNog:ENOG410PM5F,COG:U |
| chr10 | 1412654 | 1413641 | - | **hypothetical protein** | GO_component: GO:0005737 - cytoplasm [Evidence IEA],GO_function: GO:0005515 - protein binding [Evidence IEA],GO_process: | InterPro:IPR003005,InterPro:IPR004148,InterPro:IPR027267,InterPro:IPR037429,PFAM:PF03114 | note=EggNog:ENOG410QE92,COG:U |

| | | | | | | | |
|---|---|---|---|---|---|---|---|
| | | | | | GO:0007015 - actin filament organization [Evidence IEA] | | |
| chr2 | 482295 | 483598 | - | **hypothetical protein (Actin-related protein)** | | InterPro:IPR004000,InterPro:IPR020902,PFAM:PF00022 | note=EggNog:ENOG410PHDF,COG:Z |
| chr2 | 2132703 | 2133386 | + | **hypothetical protein** | GO_component: GO:0005840 - ribosome [Evidence IEA],GO_function: GO:0003735 - structural constituent of ribosome [Evidence IEA],GO_process: GO:0006412 - translation [Evidence IEA] | InterPro:IPR001047,InterPro:IPR022309,PFAM:PF01201 | note=EggNog:ENOG410PI2H,COG:J |
| chr2 | 3027726 | 3028871 | + | **hypothetical protein** | | InterPro:IPR022057,PFAM:PF12271 | note=EggNog:ENOG410PIP5,COG:S |
| chr2 | 3283071 | 3285575 | - | **hypothetical protein** | GO_component: GO:0016020 - membrane [Evidence IEA],GO_function: GO:0016491 - oxidoreductase activity [Evidence IEA],GO_function: GO:0016651 - oxidoreductase activity, acting on NAD(P)H [Evidence IEA],GO_function: GO:0051536 - iron-sulfur cluster binding [Evidence IEA],GO_function: GO:0008137 - NADH dehydrogenase (ubiquinone) activity [Evidence IEA],GO_function: GO:0009055 - electron transfer activity [Evidence IEA],GO_process: GO:0055114 - oxidation-reduction process [Evidence IEA],GO_process: GO:0042773 - ATP synthesis coupled electron transport [Evidence IEA] | InterPro:IPR000283,InterPro:IPR001041,InterPro:IPR006656,InterPro:IPR006963,InterPro:IPR010228,InterPro:IPR015405,InterPro:IPR019574,InterPro:IPR036010,PFAM:PF00111,PFAM:PF09326,PFAM:PF10588,PFAM:PF13510 | note=EggNog:ENOG410PH1J,COG:C |

| | | | | | | | |
|---|---|---|---|---|---|---|---|
| chr2 | 4291835 | 4293444 | + | **hypothetical protein** | GO_component: GO:0005737 - cytoplasm [Evidence IEA],GO_component: GO:0005852 - eukaryotic translation initiation factor 3 complex [Evidence IEA],GO_function: GO:0003743 - translation initiation factor activity [Evidence IEA] | InterPro:IPR000717,InterPro:IPR019382,PFAM:PF10255 | note=EggNog:ENOG410PFZ7,COG:J |
| chr3 | 1915050 | 1916427 | - | **hypothetical protein** | GO_function: GO:0016747 - transferase activity, transferring acyl groups other than amino-acyl groups [Evidence IEA],GO_function: GO:0003824 - catalytic activity [Evidence IEA] | InterPro:IPR002155,InterPro:IPR016039,InterPro:IPR020610,InterPro:IPR020615,InterPro:IPR020616,InterPro:IPR020617,PFAM:PF00108,PFAM:PF02803 | note=EggNog:ENOG410PG5V,COG:I |
| chr3 | 3222971 | 3225937 | - | **hypothetical protein** | GO_function: GO:0008536 - Ran GTPase binding [Evidence IEA],GO_process: GO:0006886 - intracellular protein transport [Evidence IEA] | InterPro:IPR001494,InterPro:IPR011989,InterPro:IPR016024,PFAM:PF02985,PFAM:PF03810,PFAM:PF13513 | note=EggNog:ENOG410PJ5B,COG:U,Y |
| chr3 | 3906505 | 3906918 | + | **hypothetical protein** | GO_function: GO:0046982 - protein heterodimerization activity [Evidence IEA] | InterPro:IPR003958,InterPro:IPR009072,PFAM:PF00808 | note=EggNog:ENOG410PS30 |
| chr4 | 1363062 | 1363859 | + | **hypothetical protein** | | InterPro:IPR009038,InterPro:IPR036598,PFAM:PF01105 | note=EggNog:ENOG410PK0H,COG:U |
| chr4 | 1476210 | 1476771 | - | **hypothetical protein** | GO_component: GO:0030532 - small nuclear ribonucleoprotein complex [Evidence IEA],GO_process: GO:0008380 - RNA splicing [Evidence IEA] | InterPro:IPR001163,InterPro:IPR010920,InterPro:IPR027248,PFAM:PF01423 | note=BUSCO:EOG09265IT6,EggNog:ENOG410PNRK,COG:A |
| chr4 | 2371809 | 2372304 | + | **hypothetical protein** | | InterPro:IPR005651,PFAM:PF03966 | note=BUSCO:EOG09265F2Y,EggNog:ENOG410PPKG,COG:S (Taxomomic profile: 100% Dikaryote fungi, 72 species) |

| | | | | | | | |
|---|---|---|---|---|---|---|---|
| chr4 | 2539144 | 2540372 | + | **hypothetical protein** | GO_function: GO:0005515 - protein binding [Evidence IEA] | InterPro:IPR001680,InterPro:IPR015943,InterPro:IPR017986,InterPro:IPR019775,InterPro:IPR020472,InterPro:IPR036322,PFAM:PF00400 | note=EggNog:ENOG410PJ2T,COG:A |
| chr7 | 1225597 | 1228359 | + | **hypothetical protein** | GO_function: GO:0005524 - ATP binding [Evidence IEA],GO_process: GO:0019538 - protein metabolic process [Evidence IEA] | InterPro:IPR001270,InterPro:IPR003593,InterPro:IPR003959,InterPro:IPR004176,InterPro:IPR018368,InterPro:IPR019489,InterPro:IPR027417,InterPro:IPR028299,InterPro:IPR036628,InterPro:IPR041546,PFAM:PF00004,PFAM:PF00158,PFAM:PF02861,PFAM:PF07724,PFAM:PF07728,PFAM:PF10431,PFAM:PF17871 | note=EggNog:ENOG410PGGQ,COG:O |
| chr7 | 2007804 | 2008631 | - | **hypothetical protein** | GO_component: GO:0005840 - ribosome [Evidence IEA],GO_function: GO:0003735 - structural constituent of ribosome [Evidence IEA],GO_process: GO:0006412 - translation [Evidence IEA] | InterPro:IPR021131,InterPro:IPR021132,InterPro:IPR036227,PFAM:PF17135 | note=EggNog:ENOG410PMXF,COG:J |
| chr4 | 2095406 | 2097441 | - | **Importin alpha subunit (Karyopherin alpha subunit) (Serine-rich RNA polymerase I** | GO_component: GO:0005634 - nucleus [Evidence IEA],GO_component: GO:0005737 - cytoplasm [Evidence IEA],GO_function: GO:0005515 - protein binding [Evidence IEA],GO_function: GO:0140142 - nucleocytoplasmic carrier activity [Evidence IEA],GO_function: GO:0061608 - nuclear import signal receptor activity [Evidence | InterPro:IPR000225,InterPro:IPR002652,InterPro:IPR011989,InterPro:IPR016024,InterPro:IPR024931,InterPro:IPR032413,InterPro:IPR036975,PFAM:PF00514,PFAM:PF01749,PFAM:PF02985,PFAM:PF | note=BUSCO:EOG09261OSU,EggNog:ENOG410PI6Q,COG:U |

| | | | | | | | |
|---|---|---|---|---|---|---|---|
| | | | | **suppressor protein)** | IEA],GO_process: GO:0006606 - protein import into nucleus [Evidence IEA] | 13513,PFAM:PF13646,PFAM:PF16186 | |
| chr1 | 1058610 | 1059442 | + | **kinetochore-associated Ndc80 complex subunit spc25** | | InterPro:IPR013255,PFAM:PF08234 | note=EggNog:ENOG410PHDK,COG:S |
| chr3 | 2787435 | 2788130 | + | **mitochondrial membrane protein** | GO_function: GO:0005515 - protein binding [Evidence IEA],GO_process: GO:0000266 - mitochondrial fission [Evidence IEA] | InterPro:IPR011990,InterPro:IPR016543,InterPro:IPR028058,InterPro:IPR028061,InterPro:IPR033745,PFAM:PF14852,PFAM:PF14853 | note=BUSCO:EOG09265A4E,EggNog:ENOG410PMVK,COG:M |
| chr1 | 3675964 | 3677978 | + | **Phosphoglucomutase-2** | GO_function: GO:0016868 - intramolecular transferase activity, phosphotransferases [Evidence IEA],GO_process: GO:0005975 - carbohydrate metabolic process [Evidence IEA],GO_process: GO:0071704 - organic substance metabolic process [Evidence IEA] | InterPro:IPR005841,InterPro:IPR005844,InterPro:IPR005845,InterPro:IPR005846,InterPro:IPR016055,InterPro:IPR036900,PFAM:PF02878,PFAM:PF02879,PFAM:PF02880 | note=EggNog:ENOG410PHM1,COG:G |
| chr3 | 2221373 | 2223768 | + | **Pre-mRNA-splicing factor cef1** | GO_function: GO:0003677 - DNA binding [Evidence IEA] | InterPro:IPR001005,InterPro:IPR009057,InterPro:IPR017930,InterPro:IPR021786,PFAM:PF00249,PFAM:PF11831, PFAM:PF13921 | note=BUSCO:EOG09261CXQ,EggNog:ENOG410PFBZ,COG:K |

| | | | | | | | |
|---|---|---|---|---|---|---|---|
| chr1 | 3739004 | 3739974 | - | **proteasome core particle subunit beta 2** | GO_component: GO:0005839 - proteasome core complex [Evidence IEA],GO_function: GO:0004175 - endopeptidase activity [Evidence IEA],GO_function: GO:0004298 - threonine-type endopeptidase activity [Evidence IEA],GO_process: GO:0051603 - proteolysis involved in cellular protein catabolic process [Evidence IEA] | InterPro:IPR000243,InterPro:IPR001353,InterPro:IPR023333,InterPro:IPR024689,InterPro:IPR029055,PFAM:PF00227,PFAM:PF12465 | note=BUSCO:EOG092641M3,MEROPS:MER0000542,EggNog:ENOG410PH45,COG:O |
| chr1 | 3802131 | 3802822 | - | **ribosomal 40S subunit protein S13** | GO_component: GO:0005840 - ribosome [Evidence IEA],GO_function: GO:0003735 - structural constituent of ribosome [Evidence IEA],GO_process: GO:0006412 - translation [Evidence IEA] | InterPro:IPR000589,InterPro:IPR009068,InterPro:IPR012606,InterPro:IPR023029,PFAM:PF00312,PFAM:PF08069 | note=BUSCO:EOG0926514P,EggNog:ENOG410PMVC,COG:J |
| chr2 | 2974410 | 2975180 | + | **ribosomal protein S5** | GO_component: GO:0015935 - small ribosomal subunit [Evidence IEA],GO_function: GO:0003735 - structural constituent of ribosome [Evidence IEA],GO_function: GO:0003723 - RNA binding [Evidence IEA],GO_process: GO:0006412 - translation [Evidence IEA] | InterPro:IPR000235,InterPro:IPR005716,InterPro:IPR020606,InterPro:IPR023798,InterPro:IPR036823,PFAM:PF00177 | note=EggNog:ENOG410PIAI,COG:J |
| chr1 | 3035216 | 3036146 | - | **RIBULOSE-phosphate 3-epimerase** | GO_function: GO:0004750 - ribulose-phosphate 3-epimerase activity [Evidence IEA],GO_function: GO:0016857 - racemase and epimerase activity, acting on carbohydrates and derivatives [Evidence IEA],GO_function: GO:0003824 - catalytic activity [Evidence IEA],GO_process: GO:0006098 - pentose-phosphate shunt [Evidence IEA],GO_process: GO:0005975 - carbohydrate metabolic process [Evidence IEA] | InterPro:IPR000056,InterPro:IPR011060,InterPro:IPR013785,InterPro:IPR026019,PFAM:PF00834 | note=BUSCO:EOG092644Z6,EggNog:ENOG410PGBP,COG:G |

| | | | | | | | |
|---|---|---|---|---|---|---|---|
| chr3 | 2136284 | 2136812 | + | **RNA polymerase II mediator complex subunit** | GO_component: GO:0016592 - mediator complex [Evidence IEA] | InterPro:IPR021384,InterPro:IPR037212,PFAM:PF11221 | note=BUSCO:EOG09265SHM,EggNog:ENOG410PQXU,COG:K |
| chr5 | 2053980 | 2054325 | - | **Sec61p translocation complex subunit** | GO_component: GO:0016020 - membrane [Evidence IEA],GO_function: GO:0015450 - P-P-bond-hydrolysis-driven protein transmembrane transporter activity [Evidence IEA],GO_process: GO:0006886 - intracellular protein transport [Evidence IEA],GO_process: GO:0006605 - protein targeting [Evidence IEA],GO_process: GO:0015031 - protein transport [Evidence IEA] | InterPro:IPR001901,InterPro:IPR008158,InterPro:IPR023391,PFAM:PF00584 | note=BUSCO:EOG09265PUI,EggNog:ENOG410PRYA,COG:U |
| chr5 | 2037555 | 2039619 | + | **Sulfate adenylyltransferase** | GO_function: GO:0004781 - sulfate adenylyltransferase (ATP) activity [Evidence IEA],GO_function: GO:0005524 - ATP binding [Evidence IEA],GO_function: GO:0004020 - adenylylsulfate kinase activity [Evidence IEA],GO_process: GO:0000103 - sulfate assimilation [Evidence IEA],GO_process: GO:0000096 - sulfur amino acid metabolic process [Evidence IEA] | InterPro:IPR002650,InterPro:IPR002891,InterPro:IPR014729,InterPro:IPR015947,InterPro:IPR024951,InterPro:IPR025980,InterPro:IPR027417,InterPro:IPR027535,PFAM:PF01583,PFAM:PF01747,PFAM:PF14306 | note=EggNog:ENOG410PFXE,COG:P |
| chr3 | 412800 | 415154 | + | **Translation initiation factor 3 subunit b** | GO_component: GO:0005852 - eukaryotic translation initiation factor 3 complex [Evidence IEA],GO_function: GO:0003743 - translation initiation factor activity [Evidence IEA],GO_function: GO:0003723 - RNA binding [Evidence IEA],GO_function: GO:0031369 - translation initiation factor binding [Evidence IEA],GO_function: GO:0005515 - protein binding | InterPro:IPR000504,InterPro:IPR011400,InterPro:IPR012677,InterPro:IPR013979,InterPro:IPR015943,InterPro:IPR034363,InterPro:IPR035979,PFAM:PF08662 | note=BUSCO:EOG09260K24,EggNog:ENOG410PGHS,COG:J |

| | | | | | | | |
|---|---|---|---|---|---|---|---|
| | | | | | [Evidence IEA],GO_function: GO:0003676 - nucleic acid binding [Evidence IEA],GO_process: GO:0006413 - translational initiation [Evidence IEA] | | |
| chr1 | 4128297 | 4129890 | - | **translation initiation factor eIF4A** | GO_function: GO:0005524 - ATP binding [Evidence IEA],GO_function: GO:0003676 - nucleic acid binding [Evidence IEA] | InterPro:IPR000629,InterPro:IPR001650,InterPro:IPR011545,InterPro:IPR014001,InterPro:IPR014014,InterPro:IPR027417,PFAM:PF00270,PFAM:PF00271,PFAM:PF04851 | note=EggNog:ENOG410PFHJ,COG:J |
| chr4 | 3873687 | 3877402 | + | **translational elongation factor EF-1 alpha** | GO_function: GO:0005524 - ATP binding [Evidence IEA],GO_function: GO:0016887 - ATPase activity [Evidence IEA] | InterPro:IPR003439,InterPro:IPR003593,InterPro:IPR011989,InterPro:IPR016024,InterPro:IPR017871,InterPro:IPR021133,InterPro:IPR027417,InterPro:IPR040533,PFAM:PF00005,PFAM:PF17947 | note=EggNog:ENOG410PGET,COG:J |
| chr1 | 1297736 | 1298401 | + | **Ubiquitin-conjugating enzyme E2 11** | | InterPro:IPR000608,InterPro:IPR016135,InterPro:IPR023313,PFAM:PF00179 | note=EggNog:ENOG410PNQ1,COG:O |
| chr3 | 1917440 | 1918078 | + | **Ubiquitin-conjugating enzyme E2 2** | | InterPro:IPR000608,InterPro:IPR016135,InterPro:IPR023313,PFAM:PF00179 | note=EggNog:ENOG410PHPZ,COG:K |
